# A high-quality bread wheat genome unravels adaptive evolution of wheat end-use quality

**DOI:** 10.1101/2025.07.27.667006

**Authors:** Xinyou Cao, Jijin Zhang, Yafei Guo, Yuhong Huang, Beirui Niu, Xin Gao, Jun Xu, Danping Li, Lei Guo, Xiukun Liu, Zhiliang Zhang, Lipeng Kang, Xuebing Qiu, Haosheng Li, Jianjun Liu, Baoxue Shan, Xiaoyan Duan, Congyang Yi, Yang Liu, Meng Jin, Changbin Yin, Jing Wang, Zhendong Zhao, Fei Lu

## Abstract

Understanding the genetics of superior dough performance is essential for improving wheat end-use quality. Here, we presented a *de novo* assembly of the genome of JM44, a Chinese wheat cultivar known for its exceptional end-use quality. The JM44 genome achieved reference-level quality (QV = 66.74), depicting a complete picture of complex regions containing gluten genes. Our microsynteny analysis across the *Triticum-Aegilops* complex showed that high-molecular-weight glutenin subunits (HMW-GSs) loci are highly conserved, while low-molecular-weight glutenin subunits (LMW-GSs) and α-/β-gliadins exhibited greater structural variation. These variable loci appear to have been preferentially selected by humans and contributed substantially to the evolution of wheat quality traits. Moreover, we observed that epistatic interactions between gluten genes are strong in modern cultivars, but markedly weaker in landraces, indicating the importance of epistatic selection during modern breeding. Our findings shed light on the genomics and evolution of wheat quality traits, providing valuable guidance for future breeding efforts.

## Introduction

Bread wheat (*Triticum aestivum* ssp. *aestivum*, 2*n* = 6*x* = 42, AABBDD) is a vital staple crop, providing essential sustenance to billions of people^1^. With the continued global population growth and rising living standards, the demand for high-quality wheat is increasing worldwide^2^. China, a populous country where wheat is a primary food, has been the largest wheat producer in the world since the early 1980s, currently accounting for 17.4% of global wheat production. Despite the enormous production volume, there is a persisting gap for high-quality wheat in China, leading to the importation of over 13 million tons of such wheat in 2023 (FAOSTAT, https://www.fao.org/faostat/en/#data/TCL). This gap exemplifies the pressing need to enhance wheat quality to meet evolving consumer preferences.

Wheat quality generally refers to the end-use quality or dough performance, a valuable trait during food processing that likely dates back to ancient Egypt, where sourdough bread was first invented^3^. Modern food science has established that wheat end-use quality is predominantly determined by gluten, which confers the distinctive viscoelastic properties of dough^4,5^. Gluten comprises two groups of prolamin proteins—polymeric glutenins and monomeric gliadins. Glutenins are further categorized into HMW-GSs and LMW-GSs, whereas gliadins are classified as α-/β-, γ-, and ω-types^6–8^. The type, content, and composition of these prolamin proteins largely determine the end-use quality of wheat^9–12^. Despite their importance, genomic analysis of gluten genes has rarely been reported across published wheat genomes, likely due to the complex clustering of gluten genes in the genome and highly repetitive sequences that separate them^13–26^. Consequently, the evolutionary process and adaptive mechanisms that have shaped the end-use quality of modern bread wheat remain elusive^4,27^. Addressing these challenges is crucial in advancing the breeding strategy to improve the quality of wheat.

In this study, we employed high-depth long-read sequencing to generate a high-quality genome assembly for JM44 (Jimai44), a wheat variety grown in the North China Plain, renowned for its superior end-use quality. By generating a high-resolution genetic variation map derived from bread wheat and its relatives (*n* = 485), we investigated the evolutionary dynamics of gluten genes across key stages of wheat evolution: polyploidization, domestication, Eurasian dispersal, and modern breeding. Our analysis uncovers genetic mechanisms that drive the adaptive evolution of quality traits in wheat, providing new insights into advance genetic studies and breeding of wheat end-use quality in the future.

## Results

### High-quality *de novo* genome assembly

To obtain a high-quality reference genome, we performed comprehensive sequencing of JM44 using multiple technologies (Supplementary Fig. 1 and Supplementary Table 1), including PacBio HiFi sequencing (47.68× coverage, 705.73 Gb HiFi reads)^28^, Illumina whole-genome sequencing (35.79× coverage), and high-throughput chromatin conformation capture (Hi-C, 128.96× coverage). Using the latest genome assembly methods (Supplementary Fig. 2b), we assembled these extensive datasets and generated a highly accurate reference genome of JM44 (Fig. 1a).

**Fig. 1.**
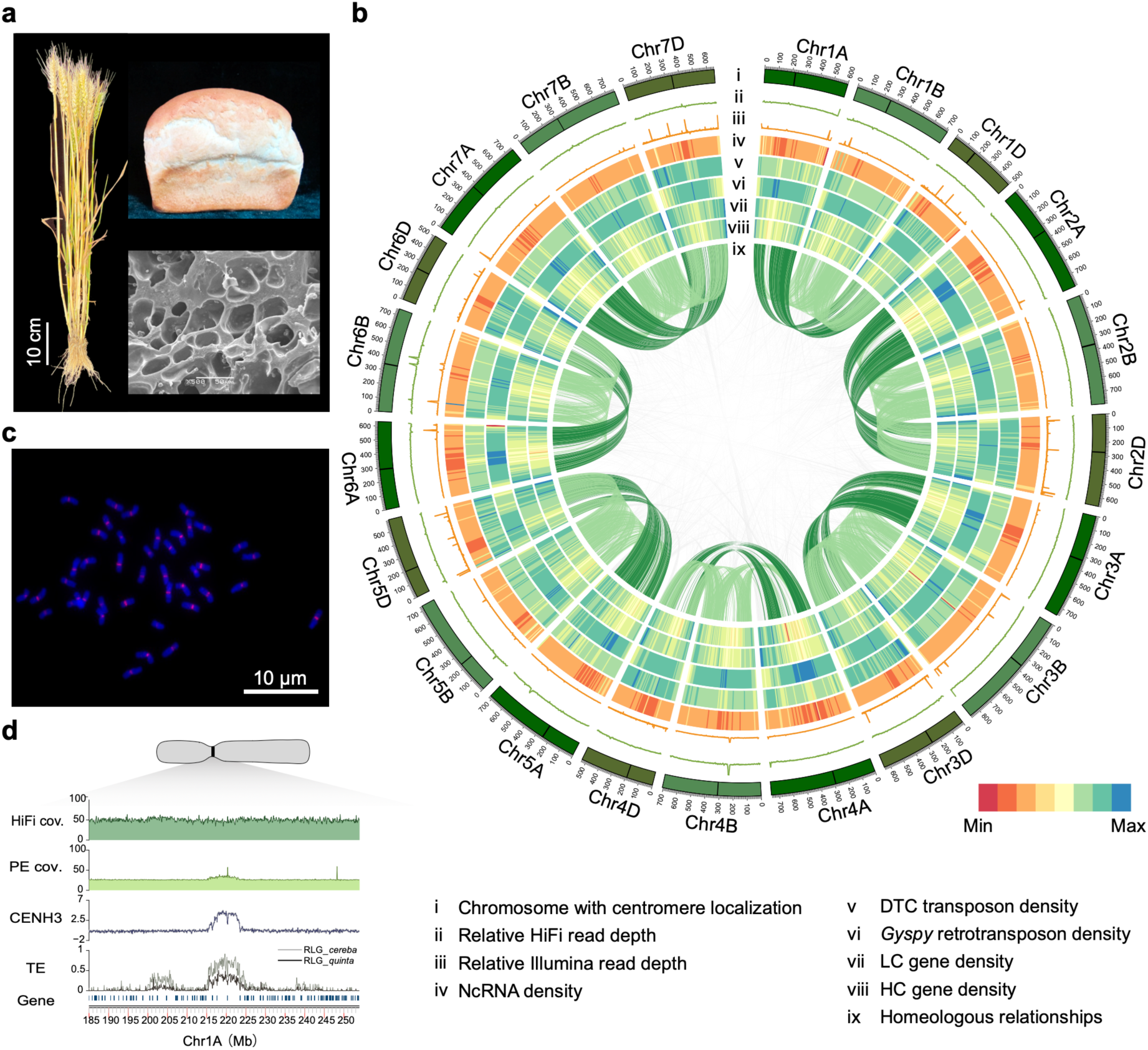
High-quality assembly and annotation of the JM44 genome. **a**, Representative images of JM44 plants, bread made from JM44 flour, and gluten microstructure. **b**, Circos plot of major features of the JM44 genome. Tracks from outer to inner circle show: (i) chromosome with centromere localization, (ii) relative HiFi read depth, (iii) relative Illumina read depth, (iv) ncRNA density, (v) DTC transposon density, (vi) *Gyspy* retrotransposon density, (vii) LC gene density, and (viii) HC gene density; the connecting lines in the center of the diagram highlight both homoeologous relationships among chromosomes and synteny between homologous genes. **c**, Fluorescence in situ hybridization of mitotic metaphase chromosomes in JM44, using CENH3 ChIP-DNA as probe (red), with chromosomes counterstained with DAPI (blue). **d**, Genomic landscape of a complete JM44 centromere. From bottom to top: (1) gene density, (2) the density of centromere-specific LTRs (RLG*_cereba* and RLG*_quinta*), (3) the density of read mapping from CENH3 ChIP-seq, (4) relative Illumina read depth; (5) relative HiFi read depth.

The final JM44 reference genome shows high continuity, with a contig N50 of 43.05 Mb and a scaffold N50 of 721.15 Mb, respectively (Supplementary Table 2). Remarkably, 14.44 Gb of the genome, accounting for 99.17% of the entire assembly, were successfully anchored to 21 pseudochromosomes (Supplementary Fig. 2b). Meanwhile, the assembly exhibits high completeness. Our k-mer analysis estimated that the actual genome size of JM44 is 14.90 Gb (Supplementary Fig. 3), indicating that 97.72% of the genome is captured in the reference. Genetic regions are particularly well-represented, with 99.3% of 1,614 conserved BUSCO genes identified (Supplementary Table 2).

Increasing continuity often comes at the cost of base-level accuracy^29^. However, the JM44 genome, supported by deep coverage of HiFi reads, achieves high base accuracy. As evaluated by Merqury^30^, it attains a quality value (QV) of 66.74. This value translates to a base error rate of less than 0.0001%, which is lower than that of published Chinese Spring reference genomes and the Kariega genome (Supplementary Table 3)^13,15,17,19,21^. Additionally, the mapped reads exhibit uniform coverage across all chromosomes (Fig. 1b and Supplementary Fig. 4). For both HiFi and Illumina sequencing, 99.35% and 99.11% of the assembly fall within three standard deviations of the mean coverage, respectively. To further assess coverage uniformity, we randomly selected 1,000 sites and compared the read depth against the expected Poisson distribution. The deviations between the observed and expected depths were negligible for both HiFi and Illumina reads (Kolmogorov-Smirnov test, *P* = 0.18 and *P* = 0.61, respectively).

### Fine structure of wheat centromeres

The centromere plays a vital role in chromosome pairing during cell division^31^. Due to the abundance of repetitive sequences in centromeric regions, resolving the fine structure of centromeres in the wheat genome has long been challenging^19,21,32,33^. To precisely define the boundaries of each centromere in the JM44 genome, we performed centromeric histone H3 (CENH3) chromatin immunoprecipitation (ChIP), followed by fluorescent in situ hybridization (FISH) of ChIPed DNA on JM44 metaphase chromosomes (Fig. 1c, d and Supplementary Fig. 5). The result showed that the centromeric regions of 18 chromosomes were completely assembled in the JM44 genome, with only three gaps remaining in the centromeres of chromosomes 2A, 5B, and 7B. The 21 centromeres had an average length of 8.2 Mb, with a relative standard deviation of 0.16. Chromosome 2B contained the longest centromere of 10.8 Mb, while chromosome 7B had the shortest one of 5.5 Mb (Supplementary Table 4 and Supplementary Fig. 5). The integrity of these centromeric regions was further supported by uniform mapping of HiFi and Illumina reads (Fig. 1d and Supplementary Fig. 5). Similar to Chinese Spring^19,21,34^, the relative positions of centromeres across chromosomes varied substantially in JM44. The L/S ratios (long arm length/short arm length) ranged from a minimum of 1.04 in Chr7A to a maximum of 2.17 in Chr5B (Supplementary Table 4).

We identified two predominant centromeric retrotransposons, RLG_*cereba* (CRW) and RLG_*quinta*^35^, in the JM44 genome. In most chromosomes, the enrichment peaks of RLG_*cereba* and RLG_*quinta* align well with the centromere (Fig. 1d, Supplementary Fig. 5). Interestingly, we observed positional shift of RLG_*cereba* and RLG_*quinta* on Chr3D and Chr4D relative to the centromere position. Similar shifting events have been reported in Chinese Spring^18^ and AK58^33^ (Supplementary Fig. 5).

Besides the centromere-specific LTRs, we also identified centromeric satellite repeats. Specifically, two distinct satellite repeats, CentT566 and CentT550, with repeat unit sizes of ∼566 bp and ∼550 bp, were observed in the JM44 genome. However, they exhibited different distribution patterns between the JM44 and Chinese Spring genomes. In JM44, CentT566 was localized to the centromeres of chromosomes 1B, 5B, and 6B, as well as adjacent regions of the centromere of chromosomes 3B (Supplementary Fig. 6), whereas CentT550 was found exclusively on the centromere of chromosome 1D (Supplementary Fig. 6). In contrast, CentT566 was distributed in the centromeres across the entire B subgenome in Chinese Spring^32^. CentT550 was present on the centromeres of chromosomes 2D, 4D, 6D, and 3B. The results suggest the presence of satellite repeats tends to be lineage-specific.

### Genome annotation of JM44

Genome annotation is essential for dissecting the genetic basis of agronomic traits. Consequently, we utilized HiFi sequencing to obtain full-length transcriptomic data from a representative set of JM44 samples, including whole seedlings, as well as roots, stems, leaves, and grains at 15 days post-flowering. By synthesizing the results from *de novo* prediction, homologous gene prediction, and transcriptome-based prediction methods (Supplementary Fig. 2c), we annotated 269,249 protein-coding genes. This includes 108,249 high-confidence (HC) genes and 161,000 low-confidence (LC) genes (Supplementary Fig. 7a). Among these, 98.17% of the HC genes and 91.64% of the LC genes were functionally annotated using five databases, including NR, EggNOG, GO, KEGG, and Swissport (Supplementary Fig. 8, Supplementary Table 5). A total of 57.07% of the HC genes were expressed at levels greater than 0.5 transcripts per million (TPM) in at least one of the tissues sampled. The total length of the HC genes is 374,353,927 bp, accounting for 2.57% of the entire genome (Supplementary Table 6). Compared to those published wheat genomes, the HC genes of JM44 have similar count and average gene length (Supplementary Table 7).

Our genome annotation encompassed a diverse array of non-coding RNAs, comprising 39,598 microRNAs, 28,266 transfer RNAs, 24,327 ribosomal RNAs, 4,043 small nucleolar RNAs, and 1,126 small nuclear RNAs. In addition, we annotated 7,233,685 transposable elements (TEs), representing 87.65% of the genome. Among the type I transposons, long terminal repeat retrotransposons were dominant, specifically the RLG_*Gypsy* and RLC_*Copia* superfamilies. For type II transposons, the DTC (CACTA) superfamily was the most prominent. Collectively, these three major superfamilies account for 43.34%, 16.13%, and 15.34% of the genome, respectively (Supplementary Table 8, Supplementary Fig. 7b). This comprehensive annotation of protein-coding genes, non-coding RNAs, and TEs complete the genome landscape of JM44 (Fig. 1b).

### Genomic structure of gluten gene loci

Gluten genes are crucial for the end-use quality of wheat but exhibit complex genomic structure^4^. To elucidate the detailed structure of these critical genes, we conducted an in-depth analysis of the gluten gene loci in the JM44 genome.

#### HMW-GSs loci

The JM44 genome contains a fully assembled and accurately annotated set of HMW-GSs genes at the homoeologous *Glu-1* loci (Fig. 2a, Supplementary Fig. 9a). These loci are located on the long arms of chromosomes 1A, 1B, and 1D, each containing two HMW-GSs genes: x-type and y-type. Among the six genes, *TaGlu-hmw-1Dx* has the longest gene body of 2,546 bp, while *TaGlu-hmw-1Ay* is the shortest, with a length of 1,826 bp (Fig. 2d).

**Fig. 2.**
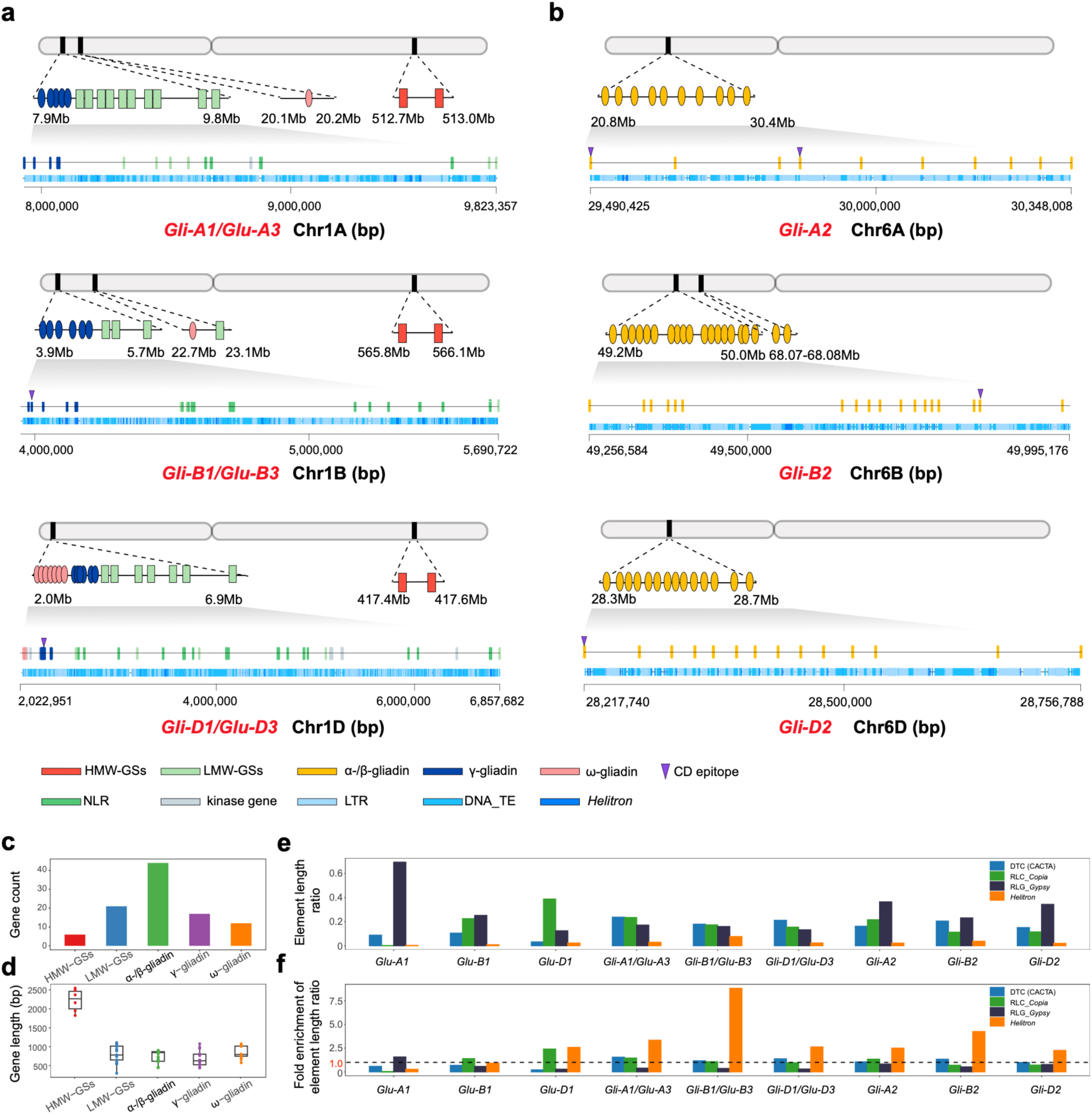
Genomic organization of gluten gene loci. **a**, Genomic organization of the *Gli-1/Glu-3* loci. The upper track shows the distribution of different gene types; the lower track shows transposable elements (TEs). Gene and TE types are color-coded, and genes containing celiac disease epitopes are marked with purple triangles. **b**, Genomic organization of the *Gli-2* loci. **c**, Count of the gluten genes. **d**, Gene length distribution of gluten genes. **e**, Proportion of total locus length occupied by major TE types within gluten loci. **f**, Fold enrichment of major TEs within gluten loci relative to the genome-wide average.

Similar to most bread wheat varieties^36^, *TaGlu-hmw-1Ay* in JM44 is silent. Earlier studies suggest that the silencing of *TaGlu-hmw-1Ay* is due to either single-nucleotide mutations that introduce premature stop codons or large LTR insertions disrupting the coding region^36^. In JM44, we identified three single nucleotide variants (SNVs) in *TaGlu-hmw-1Ay* leading to premature stop codons when compared to a normally translated *TaGlu-hmw-1Ay* gene (Supplementary Fig. 9b). However, our transcriptome analysis revealed detectable expression of *TaGlu-hmw-1Ay* in the endosperm (Supplementary Fig. 9c), suggesting that targeted editing of these SNVs could restore the activity of *TaGlu-hmw-1Ay*, and thus possibly enhance wheat quality.

Additionally, we discovered a C-to-G substitution at position 353 downstream of the *TaGlu-hmw-1Dx* transcription start site, setting JM44 apart from many other wheat varieties (Supplementary Fig. 9d). This mutation results in a serine-to-cysteine substitution, which is notable because cysteine plays a crucial role in glutenin cross-linking via disulfide bond formation, thereby impacting the elasticity of dough. We found that this additional cysteine corresponds to the superior haplotype of HMW-GS 1Dx5, corroborating previous studies on amino acids composition of gluten proteins^37^.

#### LMW-GSs and gliadin loci

For the LMW-GSs and gliadin loci, we identified 44 α-/β-gliadin genes, 17 γ-gliadin genes, 9 ω-gliadin genes, and 21 LMW-GSs genes at the *Gli-1/Glu-3* and *Gli-2* loci on the short arms of group 1 and group 6 chromosomes (Supplementary Table 9, Fig. 2a, b, and Supplementary Fig. 10). The gene lengths for LMW-GSs and gliadin range broadly from 299 bp to 1106 bp (Fig. 2d).

At the *Gli-1/Glu-3* loci, spanning approximately 2 Mb (Fig. 2a), the γ-gliadin genes are concentrated near the telomere, while the LMW-GSs genes are more diffusely distributed at the locus and interspersed with NLR and kinase genes. In contrast, ω-gliadin genes are in a compact cluster at the distal end of the *Gli-D1/Glu-D3* locus on chromosome 1D. However, on chromosomes 1A and 1B, they form a separate cluster with fewer copies, located approximately 20 Mb away from the telomere (Fig. 2a). The *Gli-2* locus spans about 0.5 Mb, consisting exclusively of α-/β-gliadin genes (Fig. 2b). There are 10, 18, and 14 α-/β-gliadin genes in the A, B, and D subgenomes, respectively. However, two α-/β-gliadin genes are found roughly 18 Mb away from the *Gli-2* locus on chromosome 6B (Fig. 2b).

Celiac disease (CD) is a common inflammatory intestinal disorder triggered by the consumption of gliadin-containing foods^4^. By searching the CD epitopes database^38^, we identified six potential CD epitopes—two in γ-gliadins and four in α-/β-gliadins (Fig. 2b, c). Targeted knockout of these epitopes could help develop wheat varieties that do not trigger human celiac disease.

#### Transposons

We conducted a detailed analysis of TEs within the gluten gene loci of *Glu-1*, *Gli-1/Glu-3*, and *Gli-2*. The results showed RLG_*Gypsy*, RLC_*Copia*, and DTC (CACTA) transposons were the most prevalent non-genic elements, accounting for 27.44%, 17.96%, and 15.18% of the sequence in these loci, respectively (Fig. 2e, Supplementary Fig. 11). The proportion of DTC (CACTA) transposons and RLC_*Copia* in gluten gene loci mirrors their genome-wide averages, with relative enrichment values of 0.99 and 1.11, respectively. In contrast, RLG_*Gypsy* was underrepresented in these loci, with a relative enrichment value of 0.62 compared to the whole genome. Notably, *Helitron* transposons were highly enriched in the gluten gene loci, with an overall relative enrichment value of 3.06. This enrichment was particularly pronounced in the *Gli-1/Glu-3* and *Gli-2* loci, peaking at 3.97 (Fig. 2e).

### Microsynteny of gluten genes

To assess the extent of conservation among gluten genes, we conducted microsynteny analyses for each gluten gene locus across the genera *Triticum* and *Aegilops*. The long-term evolutionary process has disrupted the synteny of gluten genes, as evident when comparing bread wheat with its relatives. Notably, LMW-GSs and gliadin genes showed greater copy number variation and more frequent genomic relocations than HMW-GSs genes (Fig. 3). To quantify the degree of synteny conservation, we calculated the Synteny Conservation Index (SCI) for each category of gluten genes. HMW-GSs genes exhibited a significantly higher mean value (0.90) than LMW-GSs genes (mean = 0.42, one-tailed *t*-test, *P* = 4.02 × 10^-6^), α-/β-gliadin genes (mean = 0.39, one-tailed *t*-test, *P* = 3.64 × 10^-7^), γ-gliadin genes (mean = 0.48, one-tailed *t*-test, *P* = 2.98 × 10^-5^), and ω-gliadin genes (mean = 0.17, one-tailed *t*-test, *P* = 2.86 × 10^-5^) (Supplementary Fig. 12). This finding indicates that HMW-GSs genes are more conserved than LMW-GSs and gliadin genes.

**Fig. 3.**
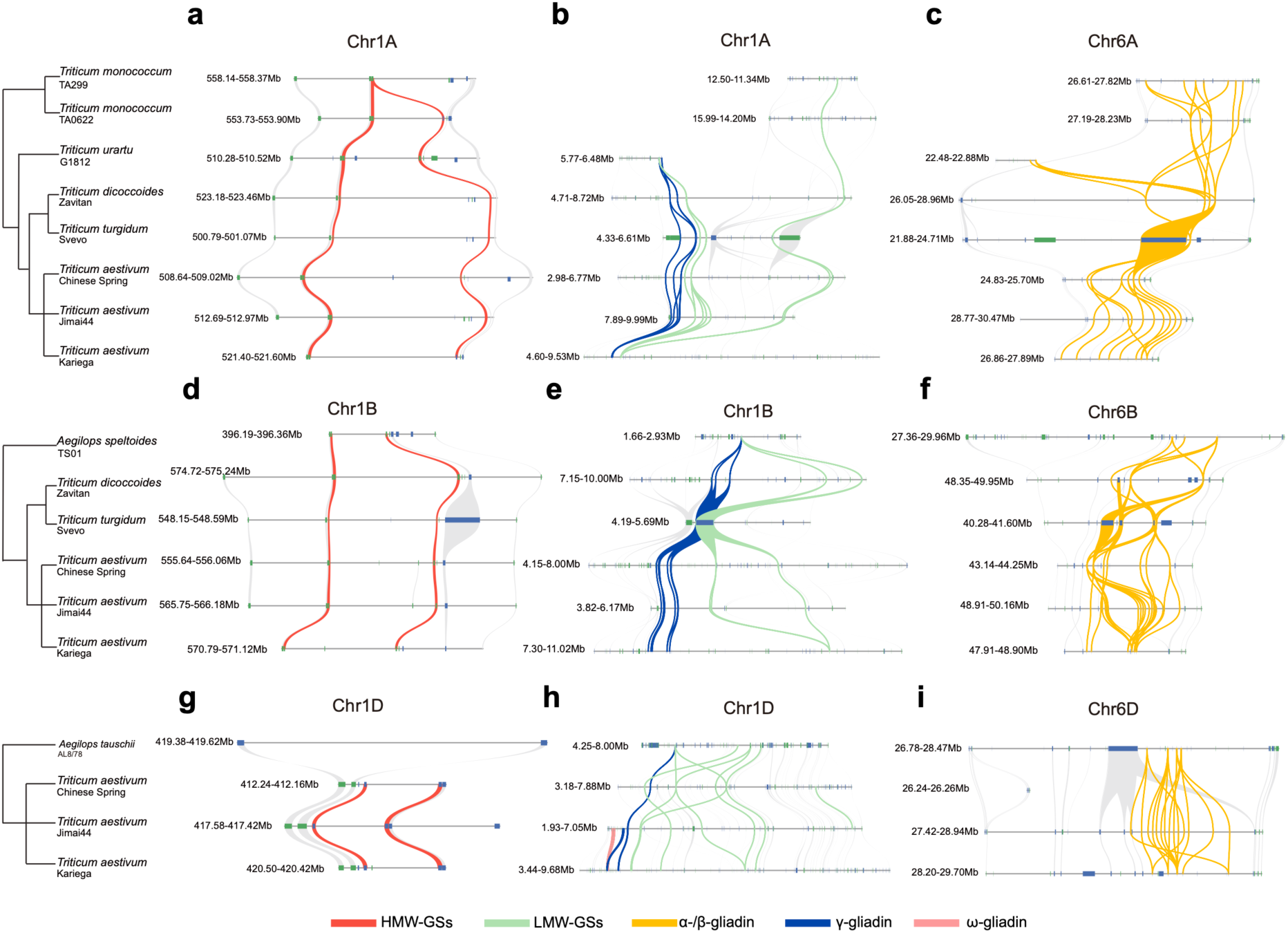
Microsynteny analysis of gluten loci across the genera *Triticum* and *Aegilops*. **a**–**c**, Microsynteny of the *Glu-1* (**a**), *Gli-1/Glu-3* (**b**), and *Gli-2* (**c**) loci in the A subgenome. **d**–**f**, Corresponding loci in the B subgenome. **g**–**i**, Corresponding loci in the D subgenome. Collinear gluten genes are connected by colored lines, while syntenic flanking genes are linked by gray lines. For the A subgenome, comparisons include wild einkorn (accession TA299), domesticated einkorn (TA10622), the A subgenome donor *Triticum urartu* (G1812), wild emmer wheat (Zavitan), durum wheat (Svevo), and three bread wheat accessions: Chinese Spring, JM44, and Kariega. For the B subgenome, comparisons include *Aegilops speltoides* (TS01), wild emmer (Zavitan), durum wheat (Svevo), and the same three bread wheat accessions: Chinese Spring, JM44, and Kariega. For the D subgenome, comparisons include the D subgenome donor *Aegilops tauschii* (AL8/78), along with Chinese Spring, JM44, and Kariega.

### Selection of gluten genes during demography events

To investigate adaptative mechanisms underlying the end-use quality of bread wheat, we constructed a high-resolution genetic variation map based on the JM44 reference genome (JVMap). To capture the key demographic events of bread wheat—domestication, polyploidization, Eurasian dispersal, and modern breeding^39^, we collected whole genome sequences of 485 accessions from bread wheat and its close relatives^22,39–45^, including bread wheat landraces (*n* = 164) and cultivars (*n* = 171), wild emmer (*Triticum turgidum* ssp. *dicoccoides*, 2*n* = 4*x* = 28, AABB, *n* = 26), domesticated emmer (*T. turgidum* ssp. *dicoccum*, 2*n* = 4*x* = 28, AABB, *n* = 29), free-threshing tetraploids (*n* = 15), and Tausch’s goatgrass (*Aegilops tauschii*, 2*n* = 2*x* = 14, DD, *n* = 62). With the representative collection, JVMap identified 86.51 million high-confidence SNPs using the cross-ploidy variation discovery pipeline we developed previously^39,41^ (Supplementary Tables 10, 11, 12, and Supplementary Note 1).

Using the cross-population composite likelihood ratio (XP-CLR) test, we detected evidence of human selection on gluten genes during the aforementioned demographic events (Supplementary Tables 14). Specifically, one, six, six, and 14 gluten genes were under selection during the four demographic events, indicating that the selection on gluten genes has been a continuous process. Interestingly, the gene *TaGlu-lmw-B3* exhibited a selection signal during the early domestication from wild emmer to domesticated emmer. Considering that leavened bread originated from ancient Egypt^3^, the result suggests that selection for end-use quality may have occurred earlier than previously assumed. Notably, *TaGlu-lmw-D2*, *TaGli-γ-D3*, and *TaGli-γ-D4*, four gluten genes that were selected during the Eurasian dispersal, were also targets of selection in modern Chinese breeding programs. This repeated selection suggests that certain gluten genes play a particularly important role in determining the end-use quality of wheat.

### Evolutionary trajectory of strong-gluten haplotypes

To elucidate the evolutionary process of end-use quality traits, we developed a haplotype inference approach that targets strong-gluten alleles, and successfully identified strong-gluten haplotypes for 19 of the 21 selected genes in the four demographic events (Supplementary Fig. 15, Supplementary Table 15, and Supplementary Note 2). For gene *TaGlu-lmw-B3*, which was under selection during the domestication stage, the strong-gluten haplotype could not be inferred due to the haplotype heterogeneity across strong-gluten varieties. Following polyploidization, six strong-gluten haplotypes, corresponding to three α-/β-gliadin genes (*TaGli-α-/β-D1*, *TaGli-α-/β-D4*, and *TaGli-α-/β-D13*) and three LMW-GSs genes (*TaGlu-lmw-B4*, *TaGlu-lmw-D4*, and *TaGlu-lmw-D5*), exhibited substantial frequency increase of superior alleles, averaging 74.34% and 67.37%, respectively. Subgenome analysis revealed that one gene from the AB subgenomes and five from the D subgenome showed increased frequencies of strong-gluten haplotypes, with average increases of 54.85% and 74.06%, respectively (Fig. 4b). These findings indicate that post-polyploidization selection significantly enhanced wheat end-use quality through the accumulation of strong-gluten haplotypes of α-/β-gliadin and LMW-GSs genes.

**Fig. 4.**
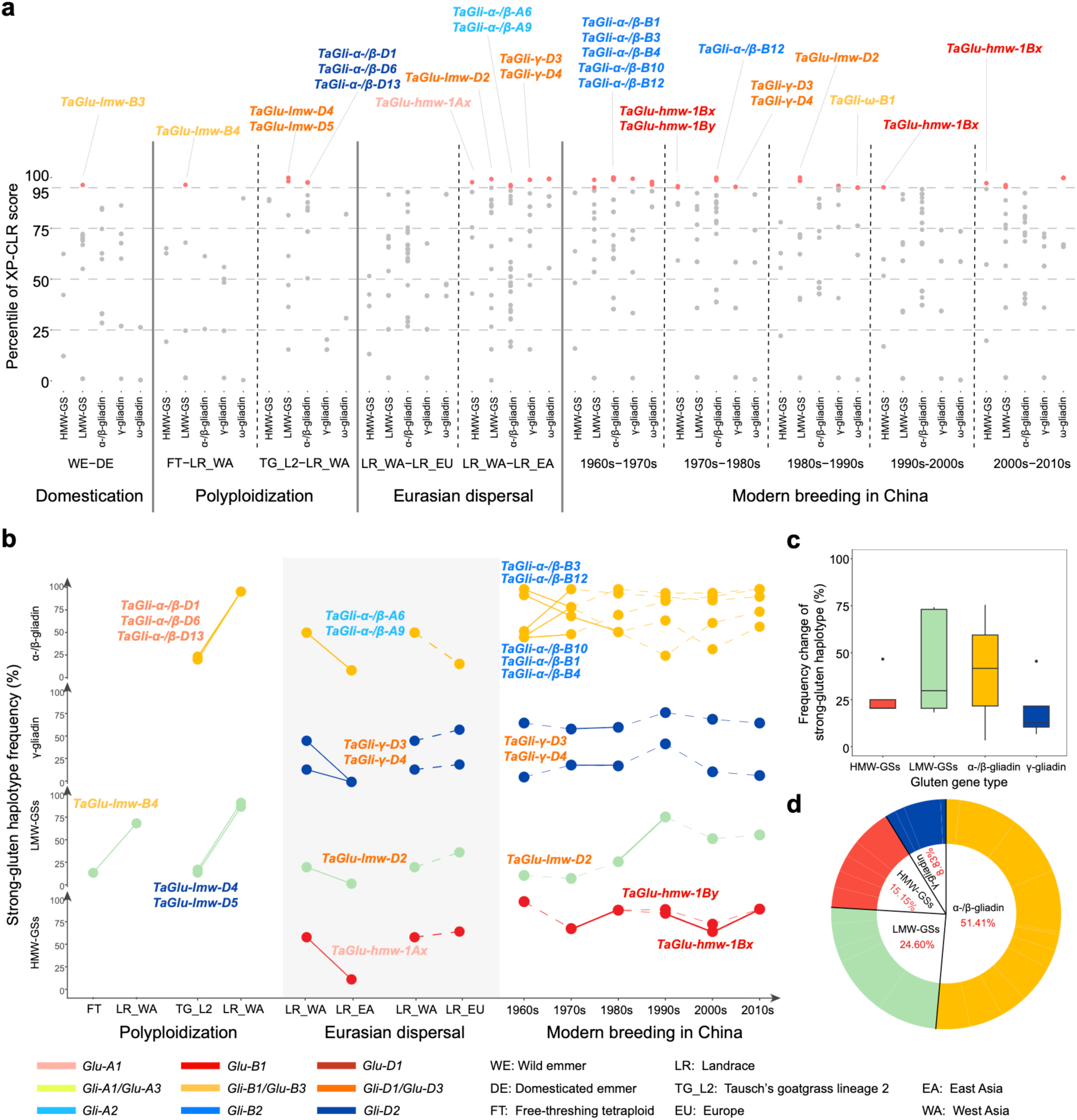
Selective sweeps and haplotype dynamics of gluten genes. **a**, Percentile of XP-CLR scores for gluten genes during key evolutionary events, including domestication, polyploidization, landrace Eurasian dispersal, and modern cultivar breeding in China. Each point represents a gluten gene, with red dots indicating those in the top 5% (putatively under selection) and gray dots representing the remainder. **b**, Dynamics of strong-gluten haplotype frequencies in selected gluten genes across distinct evolutionary stages. Solid lines denote that the gene was selected at a given stage (XP-CLR score within the top 5% significance threshold), dashed lines denote that the selection of gene did not exceed the top 5% significance threshold. **c**, Magnitude of frequency shifts in strong-gluten haplotypes among selected gluten genes, grouped by gene type. **d**, Relative contribution of each gluten gene type to the total amplitude of strong-gluten haplotype frequency changes. Abbreviations in the figure are as follows: WE, wild emmer; DE, domesticated emmer; DR, durum wheat; FT, free-threshing tetraploids; LR, landrace; WA, West Asia; EU, Europe; EA, East Asia; TG_L2, Tausch’s goatgrass lineage 2.

During the dispersal of wheat landraces from West Asia to East Asia, six glutenin genes (*TaGlu-hmw-A2*, *TaGlu-lmw-D2*, *TaGli-α-/β-A6*, *TaGli-α-/β-A9*, *TaGli-γ-D3*, and *TaGli-γ-D4*) were selected, with the frequencies of their strong-gluten haplotypes decreasing by an average of 29.75% (Fig. 4b). In contrast, during the westward spread from West Asia to Europe, most of these genes exhibited a slight increase in strong-gluten haplotype frequencies. These results showed opposing selection pressure during the eastward and westward dispersal of wheat landraces, suggesting that strong gluten quality was not favored during the ancient spread of bread wheat into East Asia.

In the stage of modern breeding in China, a total of 11 genes were selected at least once in each decade from the 1960s to the 2010s, according to our XP-CLR analysis. These selected gluten genes generally showed an increase in the frequency of strong-gluten haplotypes, with an average rise of 15.31% (Fig. 4b). However, only one gene, *TaGli-α-/β-B4*, exhibited a constant increase in the frequency of the strong-gluten haplotype, while the frequencies of other genes fluctuated over time. As HMW-GSs genes are of particular interest in modern marker-assisted breeding for end-use quality, we examined all HMW-GSs genes, regardless of whether they met the significance threshold of XP-CLR tests. We found that the superior haplotypes of *TaGlu-hmw-1Dx* and *TaGlu-hmw-1Dy*, known in breeding as the “Dx5+Dy10” combination^2^, steadily increased from 0% to 25.00% over the past 50 years (Supplementary Fig. 16a), consistent with recent breeding efforts to improve end-use quality using “Dx5+Dy10.” Although strong-gluten haplotypes of a few gluten genes showed notable increases, the overall selection efficacy on gluten genes appeared to lessen over time. Using genome-wide allele frequency changes as a baseline, we analyzed allele frequency shifts at all gluten gene loci. The results showed that allele frequency changes in gluten genes declined rapidly after the 1980s and then gradually stabilized (Supplementary Fig. 16b), which is probably due to the reduced genetic diversity among elite cultivars and insufficient functional markers for many other gluten genes^46,47^.

Considering the three stages together, we found that the frequencies of strong-gluten haplotypes in α-/β-gliadin and LMW-GSs genes changed more substantially than those of other gluten gene categories (Fig. 4c). These two gene families accounted for the majority of the total haplotype frequency shifts, contributing 51.41% and 24.60%, respectively (Fig. 4d). At the subgenome level, frequency changes of strong-gluten haplotype were more pronounced in the D subgenome than in the A and B subgenomes (Supplementary Fig. 17a), with D-subgenome gluten genes alone contributing 56.55% of the total haplotype frequency shifts (Supplementary Fig. 17b). These findings suggest that α-/β-gliadin and LMW-GSs genes have long been the key targets of quality improvement in wheat, and that the D subgenome plays a vital role in shaping of end-use quality traits in modern bread wheat.

### Epistasis between gluten genes

As wheat gluten consists of a large number and diverse types of glutenins and gliadins, we hypothesized that epistatic interactions may occur among these gluten genes, potentially contributing to the end-use quality of wheat. To test this hypothesis, we quantified the strength of epistasis between gluten genes by analyzing inter-chromosomal linkage disequilibrium (LD) among gene pairs. If epistatic interactions exist, natural selection would maintain higher LD between interacting genes than between randomly loci. Using the top 5% of LD values between random inter-chromosomal gene pairs as a threshold, we observed widespread epistatic interactions among gluten genes in both landraces and Chinese modern wheat cultivars (Fig. 5a, Supplementary Fig. 18). Notably, the overall strength of epistasis was higher in modern cultivars, with a mean LD percentile of 62.14%, compared to 46.47% in landraces (one-tailed t-test, *P* = 5.86 × 10^-43^) (Fig. 5b). Meanwhile, 79 gene pairs in the modern population exceeded the epistasis threshold, compared to only 15 pairs in the landraces (one-tailed Fisher’s exact test, *P* = 4.41 × 10^-12^) (Supplementary Note 3), indicating that epistatic interactions among gluten genes are more prevalent in modern wheat.

**Fig. 5.**
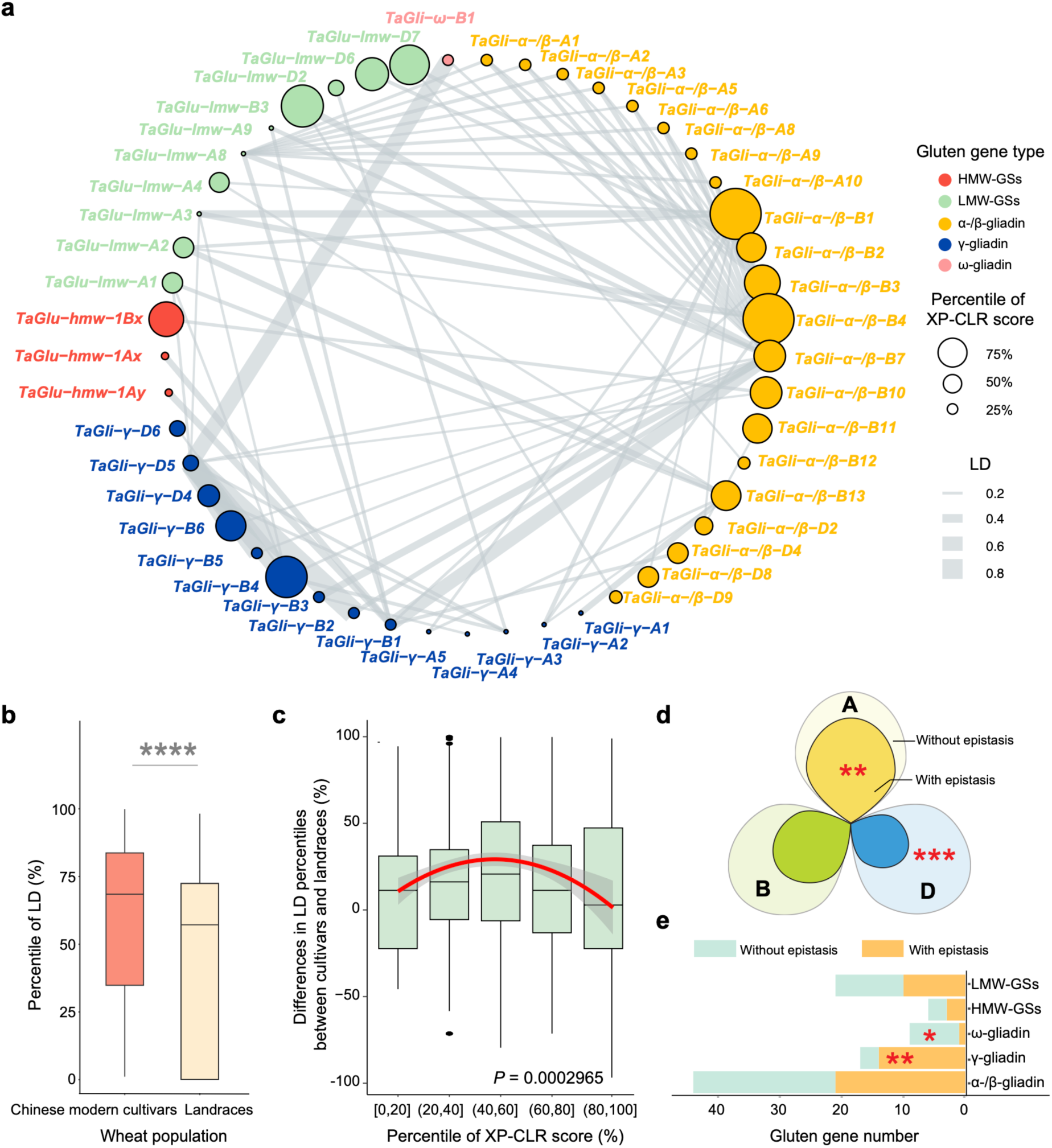
Epistatic interactions among gluten genes. **a**, Network of epistatic interactions among gluten genes in Chinese modern cultivars. Node colors represent different gluten gene families; node size indicates the strength of selection signals from landraces to modern cultivars; edge thickness reflects LD between gene pairs. **b**, Distribution of LD percentiles between gluten gene pairs in Chinese modern cultivars and landraces (one-tailed *t*-test, *P* = 5.86 × 10^-43^). **c,** Differences in LD percentiles between modern cultivars and landraces stratified by selection signal strength. *P* values for quadratic regression were calculated using two-tailed *t*-tests. **d**, Subgenome-level enrichment of gluten genes involved in significant epistasis. The A subgenome shows significant enrichment (*P* = 2.10 × 10^-3^, chi-squared test), while the D subgenome is significantly depleted (*P* = 6.00 × 10^-4^, chi-squared test). **e**, Family-level enrichment of gluten genes showing significant epistasis. γ-gliadins are significantly enriched (*P* = 3.80 × 10^-3^, chi-squared test), whereas ω-gliadins are significantly depleted (*P* = 1.31 × 10^-2^, chi-squared test).

To investigate how human selection impacts epistatic interaction between gluten genes, we examined the relationship between selection signals and the difference of LD percentile of glutens between landraces and Chinese modern cultivars. Interestingly, we observed a rise and fall of LD percentile as selection signals increased (Fig. 5c, quadratic regression, *P* = 2.97 × 10^-4^). In contrast, only a straightforward negative correlation was found between these two variables for randomly selected loci (Supplementary Fig. 20). This disparity indicates that human selection enhances epistatic interactions among gluten genes, due to their functional importance in determining the end-use quality of wheat. However, the non-linear relationship between selective sweep and epistatic strength points to complex genetic mechanisms underlying the evolution of end-use quality of wheat.

To elucidate the genetic basis underlying this non-linear relationship, we examined the distribution of epistatic interactions across subgenomes and gluten gene families. At the subgenome level, gluten genes located in the A subgenome, which harbored the fewest loci under selection for end-use quality traits, were significantly enriched for epistatic genes (chi-squared test, *P* = 2.12 × 10^-3^). In contrast, genes from the D subgenome, despite being the most frequent targets of selection, were significantly depleted for epistatic interactions (chi-squared test, *P* = 5.80 × 10^-4^) (Fig. 5d). Likewise, γ-gliadin genes, which were rarely selected during demographic events, were enriched for epistasis (chi-squared test, *P* = 3.84 × 10^-3^). Conversely, α-/β-gliadin and LMW-GSs genes were often targets for end-use quality traits. However, they were not enriched among epistatic gene pairs (Fig. 5e). These results indicate that epistatic interactions are more prevalent among gluten genes with relatively weak selection signals. Given that epistasis can create local optima in the adaptive landscape but constrain the long-term evolutionary potential^48,49^, such interactions are not generally favored by selection. Nonetheless, we did find epistatic interactions among a subset of the gluten genes, indicating that epistasis can contribute meaningful and immediate genetic gains to the end-use quality of wheat. Therefore, the epistatic architecture of gluten genes represents a promising avenue for future quality breeding efforts.

## Discussion

Meeting the global demand for high-quality wheat requires a fundamental shift in breeding strategies, prioritizing efficiency, precision, and adaptability. Evolution offers a powerful and strategic thinking—an extensive, natural field experiment that has shaped genetic architecture of traits over millennia. By decoding the molecular footprints of adaptive selection, breeding programs can move beyond empirical optimization and begin to emulate the success of evolutionary processes in tailoring wheat to diverse human needs.

The once-enigmatic and complex genomic regions—such as centromeres, telomeres, and rDNA—are now being gradually unveiled, ushering a new era of functional genome research in wheat^14,17,19,21^. In this study, we present a high-quality, chromosome-level genome assembly of the elite Chinese wheat cultivar JM44. This assembly reveals the intricate structure and composition of gluten gene loci, featuring a notable enrichment of *Helitron* transposons^50^. Furthermore, we identified mutations resulting in the silencing of *TaGlu-hmw-1Ay*^51^, providing promising targets for gene editing to improve the end-use quality of wheat.

Founded on the high-quality JM44 assembly and whole-genome sequencing data of our representative collection of wheat samples^22,39–45^, we systematically investigated evolution of all gluten genes, constructing a panoramic view of adaptive evolution for the end-use quality of wheat. By tracing the selection of gluten genes across key demographic events of bread wheat—domestication, polyploidization, Eurasian dispersal, and modern breeding in China, we found evidence of continuous human selection on end-use quality traits throughout these stages. Notably, such selection for end-use quality traits began as early as the stage of domestication. As wheat spread from West Asia across Eurasia, divergent evolution of end-use quality traits became evident, likely influenced by differing dietary cultures between the East Asia and Europe^52^. In modern Chinese breeding programs, we observed a gradual decline in selection efficacy over time, highlighting the growing need for more effective breeding strategies.

As key structural components of the gluten network, HMW-GSs form polymeric gluten webs and have long been regarded as the most important determinant of wheat end-use quality^4,53,54^. Indeed, our comparative genomic analysis revealed that HMW-GSs genes exhibit highly conserved synteny across species, reflecting their functional importance. However, HMW-GSs did not display strong selection signals during the adaptive evolution of end-use quality traits, especially in early evolutionary stages. In contrast, genes with greater genomic variability, such as LMW-GSs and α-/β-gliadins, were broadly selected and showed substantial shifts in the frequencies of strong-gluten haplotypes. The findings suggest that structural importance and genomic conservation do not inherently signify adaptive relevance, whereas genome variability may serve as an indicator of greater evolutionary adaptability. The pivotal role of LMW-GSs and α-/β-gliadins in adaptive evolution of end-use quality traits aligns well with their importance in modern breeding practices, where they are manipulated to optimize the glutenin-to-gliadin ratio and improve wheat processing quality^12,55^. Therefore, future breeding efforts should place greater emphasis on the functional value of LMW-GSs and gliadins, recognizing their genomic diversity and adaptive plasticity as key assets for further quality improvement.

Epistasis is generally disfavored by selection due to its tendency to create rugged adaptive landscapes and "mask" beneficial variations^56,57^. However, our study demonstrates that epistatic interactions among gluten genes play a pivotal role in shaping the genetic architecture of wheat end-use quality traits, with their prevalence and strength significantly amplified by human selection. Interestingly, while moderate selection signals appear to enhance epistasis, strong selection signals often diminish it, highlighting the trade-off between achieving short-term genetic gains and maintaining long-term adaptability. Notably, epistasis is enriched in loci that have been less intensively selected, such as the A subgenome and γ-gliadin genes, suggesting that these regions serve as reservoirs of epistatic genetic variation with great potential for breeding. These findings highlight that decoding and exploiting the epistatic architecture of gluten genes will be a new avenue for future quality breeding strategies.

In summary, our work provides a novel framework to understand the evolutionary process and the underlying genetic mechanisms of end-use quality traits in wheat. By integrating the demographic history of wheat and the well-recognized genetic makeup of end-use quality traits, we have revealed the adaptive history of the end-use quality trait through combined selection on evolutionarily flexible genes and epistatic interactions. These findings not only offer a theoretical foundation to understand trait evolution in wheat, but also point the way forward for future precision breeding by mimicking evolutionary processes to achieve more efficient breeding of wheat and many other crops.

## Methods

### Plants materials and genome sequencing

The hexaploid wheat cultivar JM44 were selected for genome sequencing and assembly. Seedlings of JM44 were collected for DNA sequencing and RNA sequencing. The roots, stems, leaves, embryos, and endosperms at 15 days after flowering were for RNA sequencing. For DNA sequencing, 30 single-molecule real-time cells were sequenced on PacBio Sequel II platform, and total of 705.7Gb of HiFi read was generated, and Illumina Novaseq sequencing platform was used for generating a total of 1,908.5 Gb Hi-C read dataset and 529.6 Gb paired-end dataset. RNA sequencing was performed using the PacBio Sequel II platform for the young seedlings with two single-molecule real-time cell sequencing runs. For the mixed tissues of roots, stems, leaves, embryos, and endosperms collected 15 days after flowering, one single-molecule real-time cell sequencing run was conducted. In total, approximately 20.0 GB of reads were generated. Additionally, RNA from roots, stems, leaves, embryos, and endosperms collected 15 days after flowering was subjected to TruSeq library preparation and sequenced using the Illumina NovaSeq 6000 platform, resulting in a total of 73.8GB of reads.

### Genome assembly and polishment

PacBio HiFi reads were assembled by HiFiasm (v0.14-r312)^58^ to generate initial contigs. Hi-C sequencing data were then used to hierarchically cluster and scaffold the contigs into 21 pseudochromosomes using 3D-DNA(vr180922)^59^. Manual curation of the Hi-C contact maps was performed using Juicebox (v.1.11.08)^60^. For genome polishing, both HiFi reads and Illumina paired-end reads were aligned to the assembled genome using pbmm2 (v1.9.0; https://github.com/PacificBiosciences/pbmm2) and BWA-mem2 (v2.2.1)^61^, respectively. Candidate base correction sets were generated using the HaplotypeCaller module in GATK (v4.1.9.0)^62^. Polishing edits were defined as the intersection of the two candidate sets and manually validated using IGV (v2.16.0)^63^. Finally, the consensus genome (JM44) was generated using bcftools consensus (v1.10.2-140-gc40d090)^64^, incorporating the verified polishing edits.

### Genome quality evaluation

We employed multiple complementary methods to assess the quality of the JM44 genome assembly. Assembly continuity was evaluated by calculating the contig N50 using custom shell scripts. Base-level accuracy (quality value, QV) was estimated with Merqury (v1.3)^65^ using k-mer size 21. Completeness of the gene space was assessed using the BUSCO pipeline (v5.2.2)^66^. Genome size estimation was performed by analyzing 21-mer frequency distributions using Jellyfish (v2.0)^67^ and GenomeScope2.0 (v1.0.0)^68^.

### Genome annotation

A hybrid approach involving both de novo and homology-based methods was utilized for repeat element prediction. Miniature inverted-repeat transposable elements (MITEs), a major class of type II transposons, were identified using MITE-Hunter (v1.0)^69^. Long terminal repeat (LTR) retrotransposons were detected using LTRharvest (v1.6.2)^69^ and LTR Finder (v1.07)^70^, with their predictions integrated using LTR retriever (v2.8.2)^71^. Homologous evidence was gathered using RepeatMasker (v4.0.1)^72^ to search the genome sequence against the RepBase repetitive sequence database (http://www.girinst.org/repbase) for known repetitive sequences within the target genome. By combining de novo and known repetitive sequences, RepeatMasker masked repetitive elements in the target genome. Furthermore, RepeatModeler (v2.0)^73^ was employed to de novo identify other repetitive sequences in the repeat-masked genome. Tandem repeats finder (v4.09.1)^74^ was used to detect tandem repeat. Ultimately, these diverse methods led to the extraction of repetitive sequences.

Gene structure annotation integrated ab initio prediction, homology-based inference, and transcript evidence. AUGUSTUS (v3.2.2)^75^, SNAP (v6.0)^76^, GlimmerHMM (v3.0.4)^77^, and GeneMark-ESSuite (v4.57)^78^ were utilized to predict gene structures in the repeat-masked genome. GeMoMa (v.7.1)^79^ was used for homology prediction to derive exon and intron boundary information through transcript-genome comparisons. High-quality full-length transcripts were generated from ISO-seq using SMRTLink (v11.1) (https://www.pacb.com/support/software-downloads/). PASA (vr20140417)^80^ software was utilized to predict open reading frames (ORFs) based on acquired full-length transcript sequences. The EvidenceModler^81^ integrated the aforementioned prediction outcomes, while PASA software was used to predict variable cut annotations such as UTRs. All protein-coding genes were aligned to three integrated protein sequence databases: NR (https://www.ncbi.nlm.nih.gov/protein/), SwissProt (https://ftp.uniprot.org/pub/databases/uniprot/current_release/knowledgebase/complete/uniprot_sprot.fasta.gz), eggNOG (http://eggnog5.embl.de/#/app/home). Protein domains were annotated by InterPro^82^ and the Gene Ontology^83,84^ terms for each gene were obtained from the corresponding InterPro entry. The pathways in which the genes might be involved were assigned by BLAST (v2.9.0)^85^ against the KEGG databases (https://www.genome.jp/kegg/brite.html). The protein-coding gene functional annotation results were merged from the above methods.

Non-coding RNAs were annotated using tRNAscan-SE (v2.0)^86^ for tRNAs, and BLAST searches against the Rfam database for other ncRNAs. NLR gene signatures were identified using NLR-Annotator (v2.0)^87^.

### Comparative Analysis of Homoeologous Genomes

Homoeologs were identified by detecting orthologous gene pairs within subgenomes, including subgenome-specific inparalogs resulting from post-hybridization duplications. Ortholog identification was performed using OrthoFinder (v2.5.5)^88,89^. Microsynteny of homoeologous gene pairs, defined as the conservation and collinearity of local gene order within orthologous chromosomal regions, was analyzed using MCScanX^90^.

### Gene expression analysis

The RNA sequencing data obtained from Illumina was aligned to the JM44 genome using STAR (v2.5.2a)^91^, and subsequent analysis was conducted using the EdgeR R package (v3.16.5)^92^.

### Centromere ChIP-seq data analysis

ChIP was conducted as previously described by Su^32^ with some modifications. A polyclonal antibody against wheat CENH3 was used for ChIP experiments. The specificity of these antibodies was confirmed through immunofluorescence assays on JM44’s mitotic chromosomes. Nuclei were isolated from 3-week-old seedlings and then digested using micrococcal nuclease (Sigma-Aldrich) to release nucleosomes. The digested mixture was incubated overnight with 3 μg of antibodies at 4 °C. Dynabeads Protein A (Invitrogen) were utilized to capture target antibodies from the mixture, resulting in ChIP DNA. A mock DNA control was established using input DNA under the same conditions but without antibodies. These ChIP experiments were carried out in two biological replicates. Subsequently, library construction was executed utilizing the TruSeq ChIP Sample Prep Kit (Illumina), following the manufacturer’s instructions.

The raw ChIP and input control sequencing reads underwent quality filtering and adapter sequence removal using fastp (v1.0.1)^93^. Subsequently, the trimmed reads were mapped to their respective genomes using bowtie2(v2.5.1)^94^ with default settings. The resulting file was filtered using SAMtools(v1.10.2)^95^ view to retain only reads that aligned across their entire read length. The ratio of ChIP/input coverage was determined using the deepTools(v3.5.3)^96^ function bamCompare, employing a MAPQ ≥ 30 threshold.

Fluorescence *In Situ* Hybridization was employed in this study. Root-tips of JM44 were grown at room temperature for two to three days, reaching a length of 2–3 cm. Mitotic chromosome spreads were prepared following a previously published method^3^. Fresh root-tips underwent treatment in a nitrous oxide gas chamber for 2.5 h, followed by fixation in 90% acetic acid for 10 min. ChIPed DNA was labeled with Alexa Fluor-594-5-dUTP (red).

### Centromere specific transposon analysis

Centromere-enriched retrotransposons *RLG_Cereba* and *RLG_Quinta* were identified using BLAST. Their abundance within annotated centromeric regions was quantified using Bedtools (v2.30.0)^96^.

### Microsynteny analysis of gluten loci

To identify syntenic gene blocks between JM44 and other accessions, all-against-all BLASTP (E value < 1 × 10−5, retaining the top five hits) was performed for the gene sets of each genome pair. Syntenic blocks were defined based on the presence of at least five synteny gene pairs using the MCscan package (https://github.com/tanghaibao/jcvi/wiki/MCscan-(Python-version)) with default settings. For the A subgenome, comparisons include wild einkorn (accession TA299)^97^, domesticated einkorn (TA10622)^97^, the A subgenome donor *Triticum urartu* (G1812)^98^, wild emmer wheat (Zavitan)^99^, durum wheat (Svevo)^100^, and three bread wheat accessions: Chinese Spring^13^, JM44, and Kariega^17^. For the B subgenome, comparisons include *Aegilops speltoides* (TS01)^101^, wild emmer (Zavitan), durum wheat (Svevo), and the same three bread wheat accessions: Chinese Spring, JM44, and Kariega. For the D subgenome, comparisons include the D subgenome donor *Aegilops tauschii* (AL8/78)^102^, along with Chinese Spring, JM44, and Kariega. The synteny conservation index (SCI) was used to assess the conservation of gluten genes, calculated as: SCI = Number of genes with synteny / Total number of gluten genes in JM44.

### Selective sweeps detection in different groups of wheat

Selective sweeps were identified using the cross-population composite likelihood ratio (XP-CLR) v1.0^103^ across different groups of bread wheat accessions. Wild emmer and domesticated emmer were used to investigate the domestication process of wheat. Tausch’s goatgrass, free-threshing tetraploids, and the landrace group from West Asia were used to study the polyploidization stage. Landraces from West Asia, Europe, and East Asia were employed to examine the dispersal of wheat across Eurasia. Cultivar groups from different eras in China were used to analyze the modern breeding process of wheat in China. XP-CLR scores between population pairs were calculated with parameters of --maxsnps 500 --size 100000 --step 50000. Genetic distances were based on published recombination rate data^4^. Genomic regions in the top 5% of XP-CLR scores were considered putative targets of selection. To validate selective signals, haplotype analysis was performed on gluten genes located within identified sweep regions. Gluten genes showing a haplotype frequency shift >10% were retained as high-confidence targets under selection.

### Inference of strong-gluten haplotypes

To define strong-gluten haplotypes, we used genotype data from high-quality published wheat genomes with known strong-gluten properties, including CDC Landmark, CDC Stanley, Jagger, LongReach Lancer, Mace, and JM44. Gluten gene sequences from these accessions were aligned to construct a haplotype reference set for each gluten gene. Haplotypes that were predominant among these strong-gluten varieties were designated as strong-gluten haplotypes.

### Epistatic association analysis among gluten genes

Inter-chromosomal linkage disequilibrium (LD) was calculated for 10,000 randomly selected SNPs within gene regions in both landraces and modern Chinese cultivars using PLINK (v1.90b6.21)^104^. These analyses generated background LD distributions for each population. The 95th percentile of each distribution was used as the significance threshold; LD values exceeding this threshold were considered indicative of putative epistatic interactions, thereby minimizing the influence of random associations and population structure. Using this approach, inter-chromosomal LD was specifically assessed for gluten gene pairs in both populations. Only gene pairs with LD values surpassing the population-specific threshold were retained as significant epistatic interactions.

## Data availability

All raw sequencing data, assembly results of JM44 genome generated in this study have been deposited in the Genome Sequence Archive (https://ngdc.cncb.ac.cn/gsa/) under accession number PRJCA020806. Genome annotation files are available at https://github.com/zhangjijin/JM44_genome.

## Acknowledgements

This work was supported by the National Key Research and Development Program (2021YFF1000203), the National Natural Science Foundation of China (32225038 and 32270269), National Key Research and Development Program (2022YFF1002904 and 2023YFF1000604), the Hainan Yazhou Bay Seed Lab (B21HJ0001 and B21HJ0111), the Taishan Scholars Program (tstp20240843), the Agriculture Research System of China (CARS-03-06), and the Natural Science Foundation Project of Shandong Province (ZR2022MC155).

## Author contributions

F.L. and Z.Z. designed and supervised the research. X.C., J.Z. and F.L. developed the general schematic workflow and performed data analysis. Y.G., J.X., Z.Z., L.K., B.N., J.M., X.D, B.S., and C.Y. helped with data analysis. X.L., X.Q, H.L., D.L., and L.G. helped the image acquisition. Y.H. and F.H. performed ChIP experiments. Z.Z., X.G., and J.L. make suggestions for the manuscript. C.Y. and J.W. collected plant material information. X.C., J.Z., and F.L. wrote the manuscript. All authors discussed the results and commented on the manuscript.

## Competing interests

The authors declare no competing interests.

## Notes

### Competing Interest Statement

The authors have declared no competing interest.

