## Supplementary information for "A high-quality bread wheat genome unravels adaptive evolution of wheat end-use quality"

**This Supplementary Information file includes:**

Supplementary Notes 1-3, which contain additional information on the methodology, results.

Supplementary Figs. 1-20, which present additional results and quality checks.

Supplementary References.

### Supplementary Note 1

**Construction of JM44 genome-based variation map.** To improve the accuracy of subsequent analyses, we constructed a whole-genome variation map using the JM44 genome as the reference. This map encompasses representative samples from key stages in wheat evolution, including: (i) the polyploidization event that gave rise to hexaploid wheat through hybridization between emmer wheat and Tausch's goatgrass; (ii) the domestication of emmer wheat; (iii) the global dispersal of bread wheat landraces; and (iv) the modern breeding of cultivars in China (Supplementary Fig. 13).

Specifically, the map includes 62 accessions of Tausch's goatgrass, comprising 21 from lineage 1 (L1), 30 from lineage 2 (L2), and 11 from lineage 3 (L3)[1]. It also includes 88 tetraploid wheat accessions—26 wild emmer, 29 domesticated emmer, 18 durum wheat, and 15 other free-threshing type[2–4]s. For hexaploid wheat, the dataset contains 164 landraces (22 from West Asia, 24 from East Asia, and 26 from Europe) and 171 modern cultivars, of which 105 are from China, representing distinct breeding eras[2–7]. Among the Chinese cultivars, 20th-century materials include 15 accessions from the 1960s, 10 from the 1970s, 21 from the 1980s, and 23 from the 1990s, while 21st-century materials include 24 accessions from the 2000s and 12 from the 2010s (Supplementary Tables 10, 11, 12).

Using a cross-ploidy variation discovery pipeline[2,3], we identified approximately 86 million high-quality single nucleotide polymorphisms (SNPs) and constructed JVMap, the first version of a whole-genome genetic variation map for wheat, based on the high-quality JM44 reference genome. Principal component analysis (PCA) effectively differentiates between the subgroups of tetraploid and hexaploid wheat, thereby indirectly validating the reliability of the SNP dataset (Supplementary Fig. 14a, b). On average, JVMap detects one SNP every 129 bp in the A and B subgenomes and one every 1,111 bp in the D subgenome (Supplementary Fig. 14c; Supplementary Table 13). The false-positive rate for variant calling—measured as the proportion of segregating sites in the reference accession JM44—was only 0.0056%, which is lower than previously reported rates for high-quality SNP datasets[1–8]. JVMap provides substantially improved resolution, particularly at gluten gene loci, where it identifies 751 SNPs (Supplementary Fig. 14d). As the first wheat genomic variation map with both high accuracy and fine-scale resolution constructed from a high-quality reference genome, JVMap effectively captures genetic diversity across representative populations that span all major stages of wheat evolution—including polyploidization, domestication, landrace Eurasian dispersal, and modern breeding in China.

#### Supplementary Note 2

**Inference of strong-gluten haplotypes of gluten genes.** To further investigate how strong-gluten haplotypes of gluten genes were selected during the adaptive evolution of wheat quality trait, we analyzed changes in the frequency of these strong-gluten haplotypes. Due to the lack of prior knowledge about strong-gluten haplotypes for most gluten genes, we developed an inference method to identify strong-gluten haplotypes of the selected gluten genes. The core of the inference method is that strong-gluten variety-major haplotypes are identified as the strong-gluten haplotypes (Supplementary Fig. 14) We used experimentally validated gluten genes (*TaGlu-hmw-A1*, *TaGlu-hmw-B1*, *TaGlu-hmw-B2*, *TaGlu-hmw-D1*, *TaGlu-hmw-D2, TaGli-γ-B1*, *TaGli-γ-D6*)[9–11]. with known strong-gluten haplotypes from published studies as a test set to evaluate the accuracy of our inference method, which achieved 100% accuracy. Using this approach, we inferred the strong-gluten haplotypes of 19 selected gluten genes (Supplementary Table 15), providing a basis for further investigation of their evolutionary trajectories and contributions to wheat end-use quality.

#### Supplementary Note 3

**Testing epistatic interactions among gluten genes.** We randomly selected 10,000 SNPs located within genic regions from both the landrace population and the modern Chinese cultivar population to calculate LD across chromosome. These LD served as the background distribution for testing epistasis. Using the top 5% of this distribution as a right-tailed threshold, we considered gene pairs with correlation values exceeding this threshold to exhibit significant epistatic interactions. The thresholds were 0.084 for the landrace population and 0.168 for Chinese modern cultivars (Supplementary Fig. 19). Based on these thresholds, 79 and 15 gene pairs were identified as significantly epistatic in the modern cultivar and landrace populations, respectively. To evaluate the statistical significance of this difference, we performed random pairing of gluten genes on each chromosome according to their chromosomal distribution, resulting in a total of 3,878 possible gene pairs. Using this as the denominator, a proportion test revealed that the number of epistatic gene pairs in the modern cultivars was significantly higher than that in the landraces (one-tailed Fisher's exact test, $P = 4.41 \times 10^{-12}$).

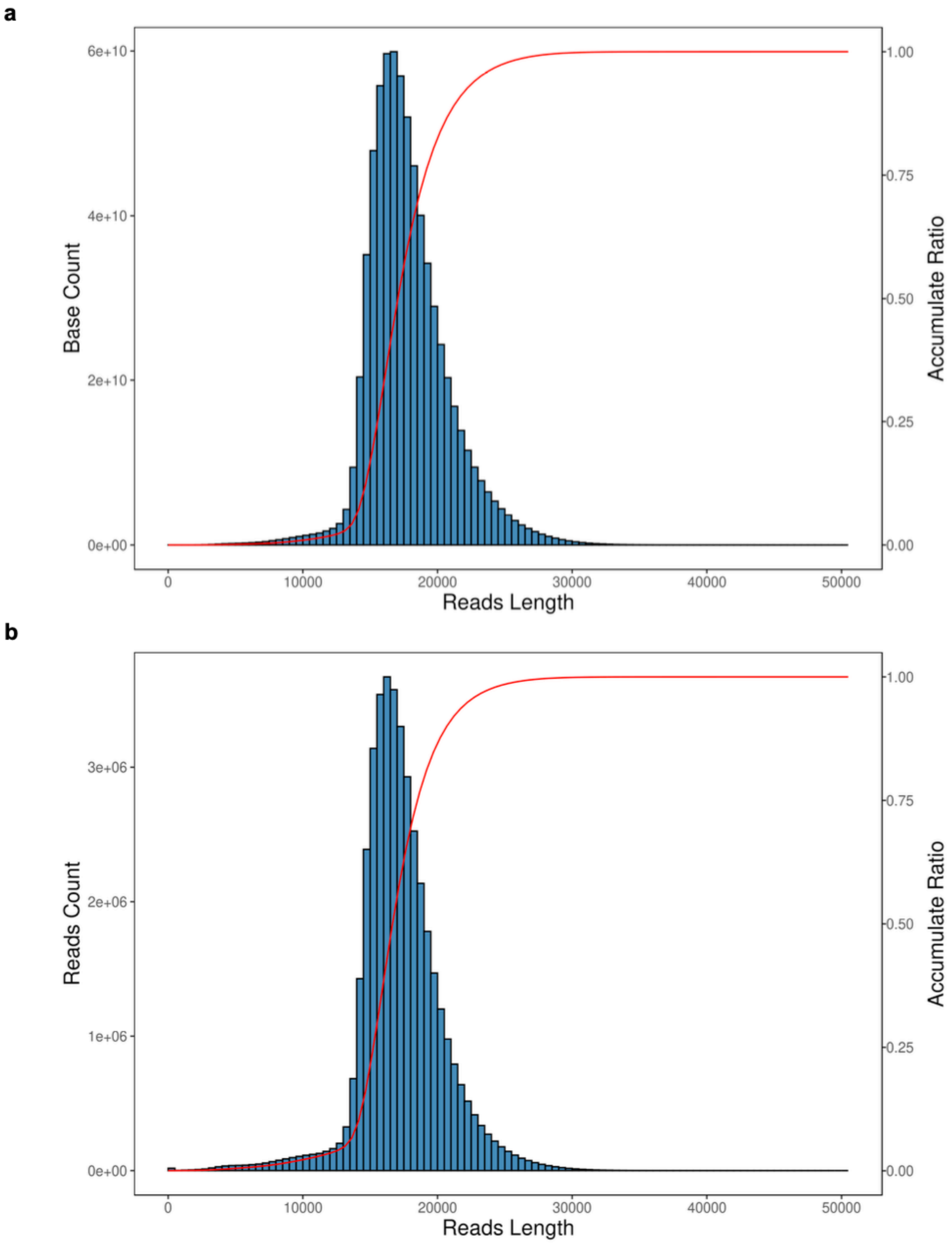
a
Base Count
6e+10
4e+10
2e+10
0e+00
Accumulate Ratio
1.00
0.75
0.50
0.25
0.00
0
10000
20000
30000
40000
50000
Reads Length
b
Reads Count
3e+06
2e+06
1e+06
0e+00
Accumulate Ratio
1.00
0.75
0.50
0.25
0.00
0
10000
20000
30000
40000
50000
Reads Length

**Supplementary Fig. 1. Summary of HiFi reads in this study. a**, Number of bases across different sequence lengths. The bar plot shows the distribution, and the red line represents the cumulative distribution. **b**, Number of sequences across different sequence lengths.

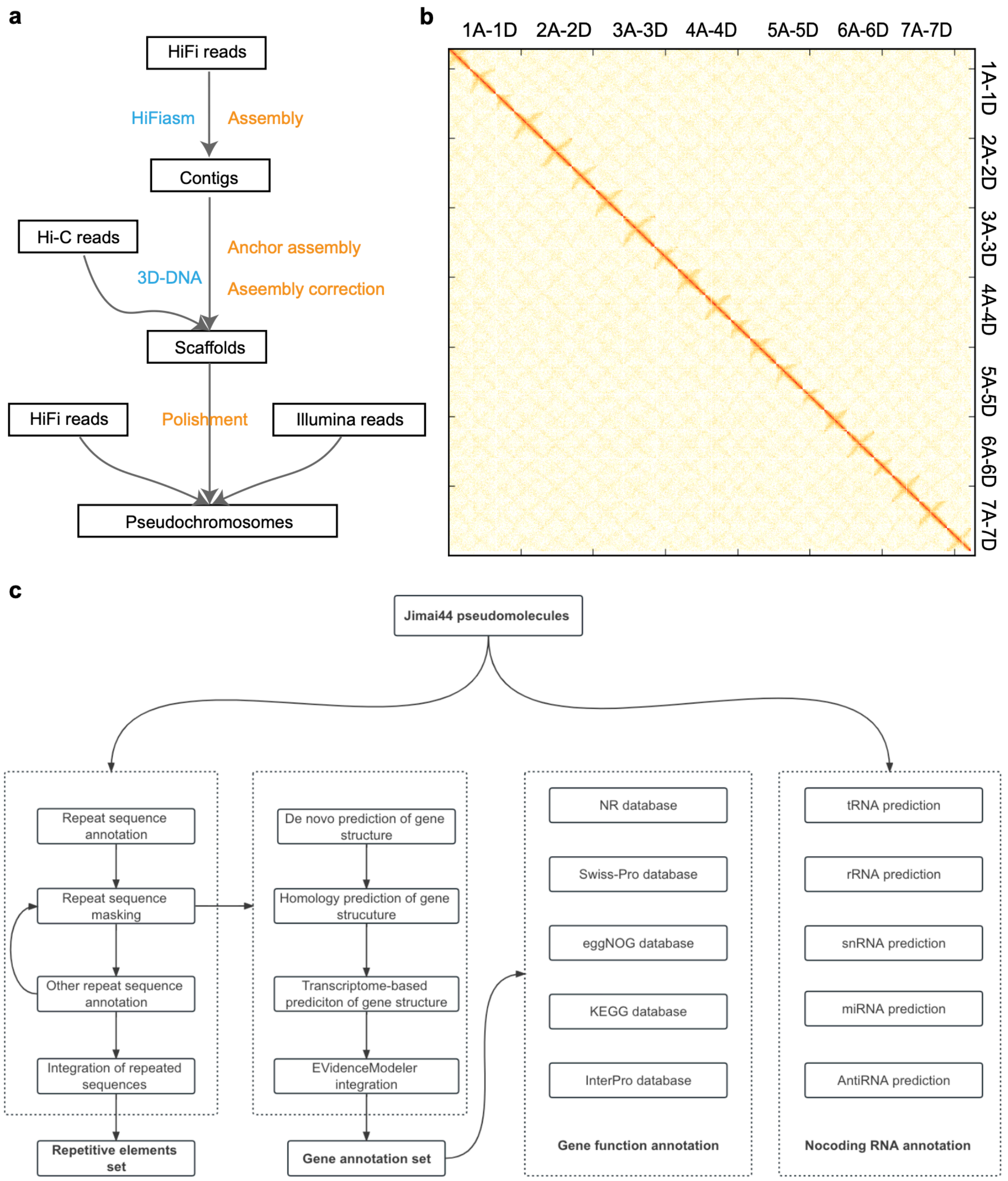


**Supplementary Fig. 2. Genome assembly and annotation strategy. a**, The process of genome sequencing and assembling of JM44. **b**, Assignment of hybrid scaffolds on chromosomes with Hi-C. **c**, The process of genome annotation of the JM44 genome.

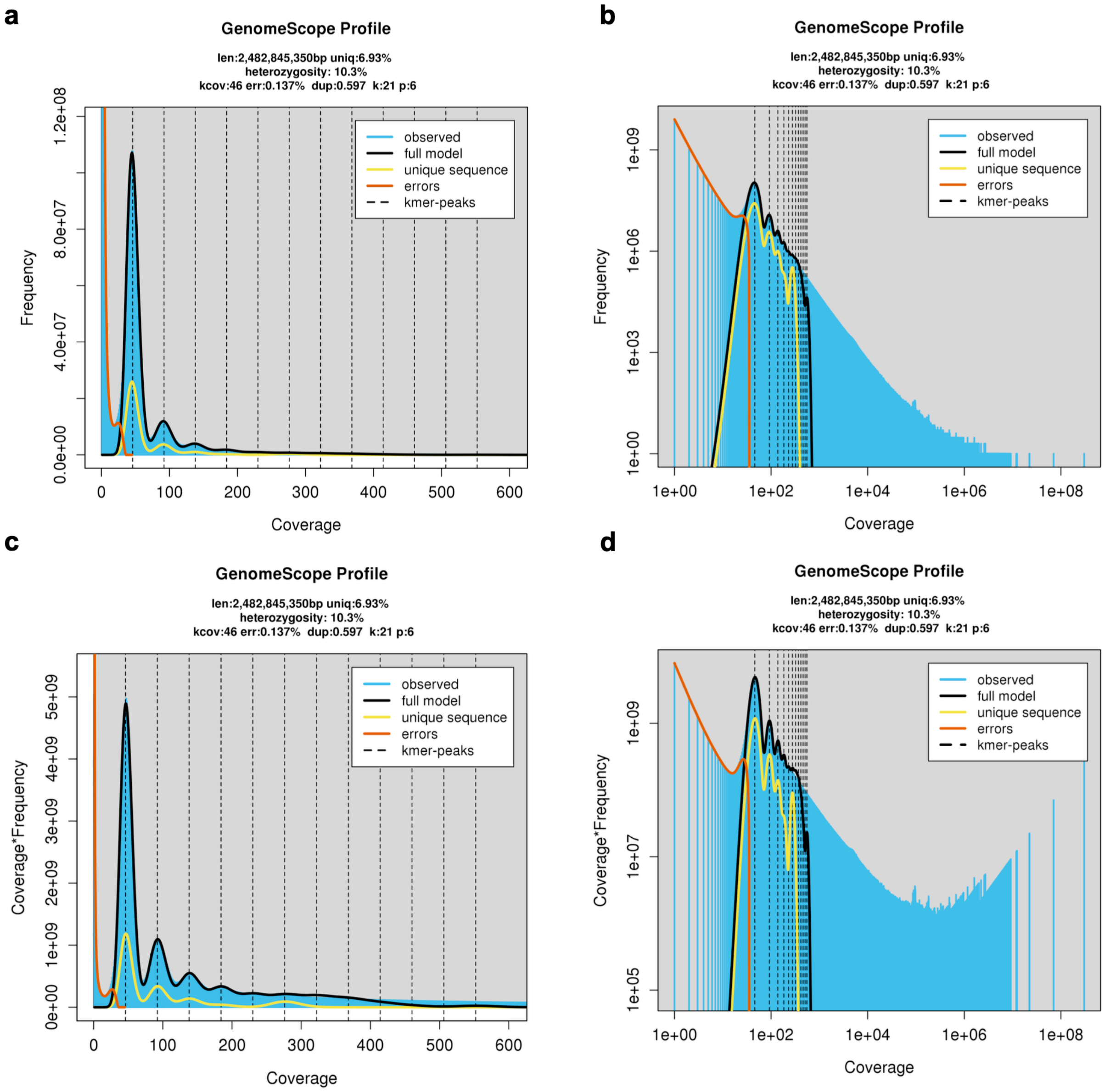


**Supplementary Fig. 3. GenomeScope results of JM44. Plots of the best fit model overlaying the k-mer spectrum for: a** untransformed linear, **b** untransformed log, **c** transformed linear, and **d** transformed log.

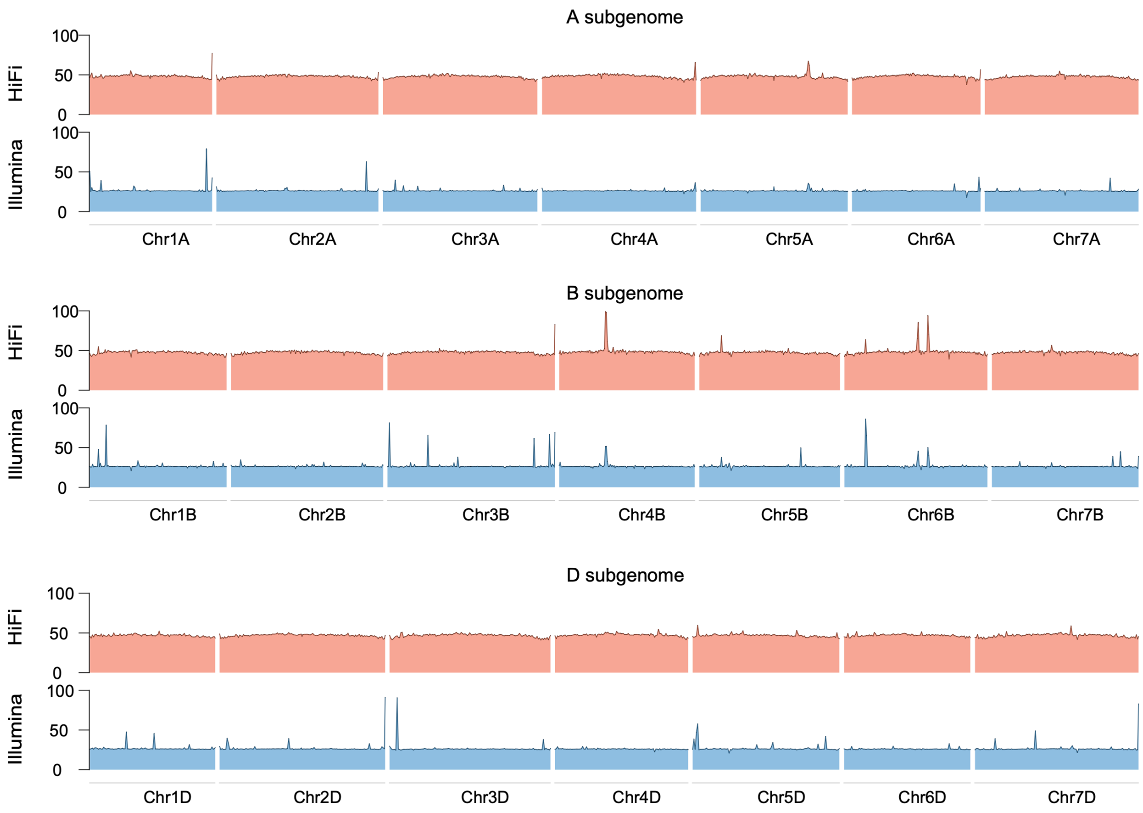


**Supplementary Fig. 4. Depth distribution of JM44 sequencing data re-mapped to genome.** The genome-wide coverage of HiFi and Illumina reads mapped are in the first and second layers from top to bottom, respectively. A convex upward peak indicates an incomplete assembly, and a downward recessed peak indicates an assembly error.

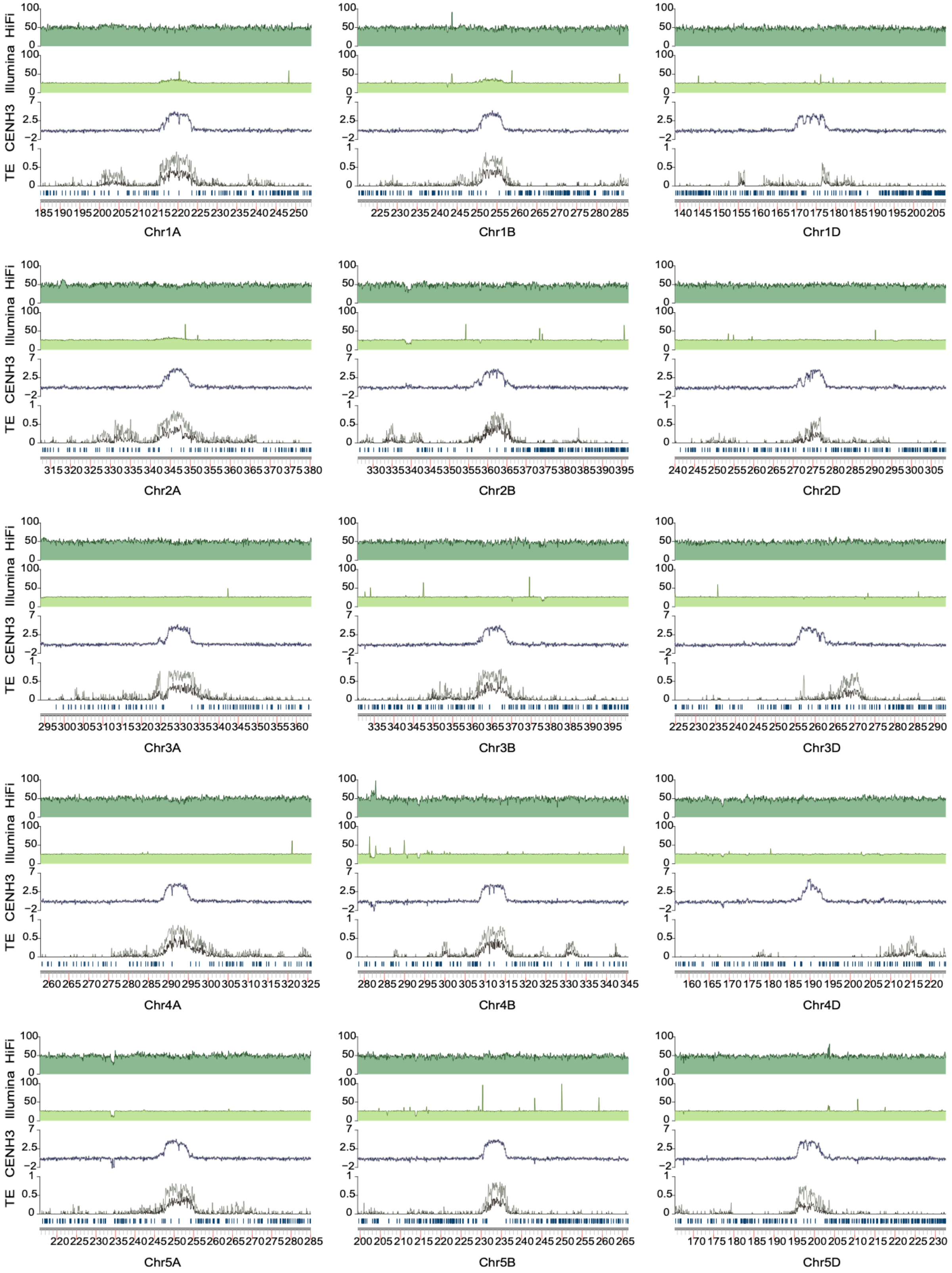
HiFi
Illumina
CENH3
TE
Chr1A
Chr1B
Chr1D
Chr2A
Chr2B
Chr2D
Chr3A
Chr3B
Chr3D
Chr4A
Chr4B
Chr4D
Chr5A
Chr5B
Chr5D

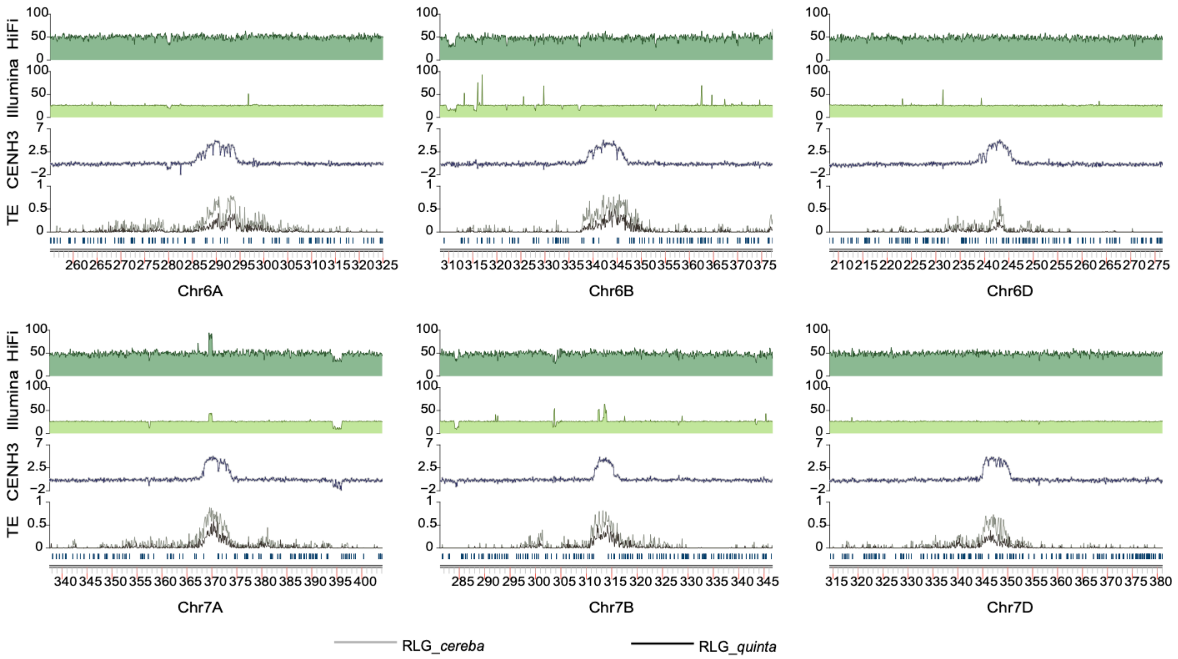


**Supplementary Fig. 5. Characteristics of the complete centromere of JM44.** The layers of every graph from bottom to top indicate (1) gene density; (2) the density of centromere specific LTRs (RLG_*cereba* and RLG_*quinta*); (3) the density of read mapping from CENH3 ChIP-seq; (4) the density of resequencing pair-end read mapping; (5) the density of resequencing HiFi read mapping.

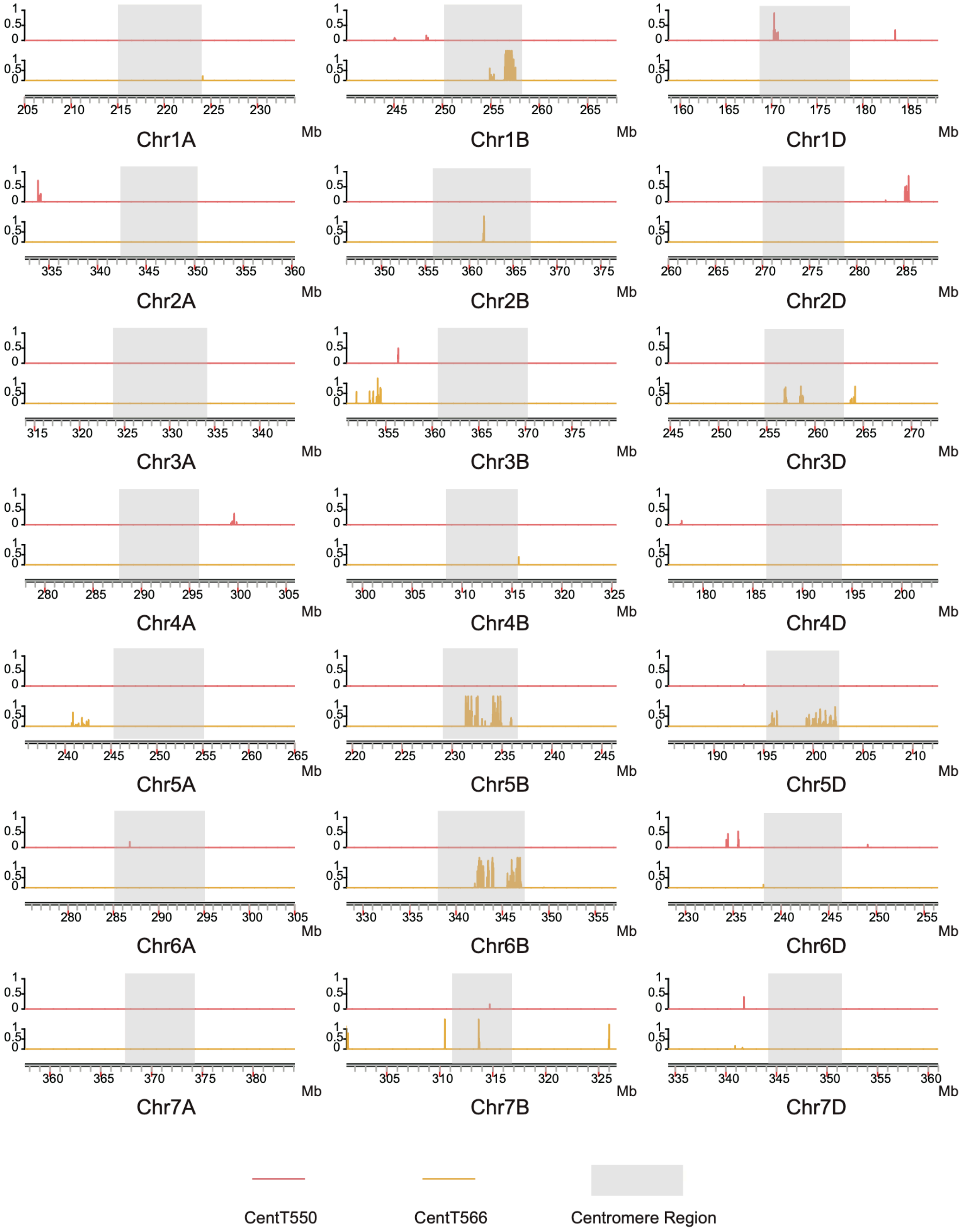


**Supplementary Fig. 6. Distribution of JM44 centromere-specific satellite repeats across centromeric regions.** The y-axis represents the density of satellite repeats within 1-kb genomic windows. Red indicates CentT550, and yellow indicates CentT566.

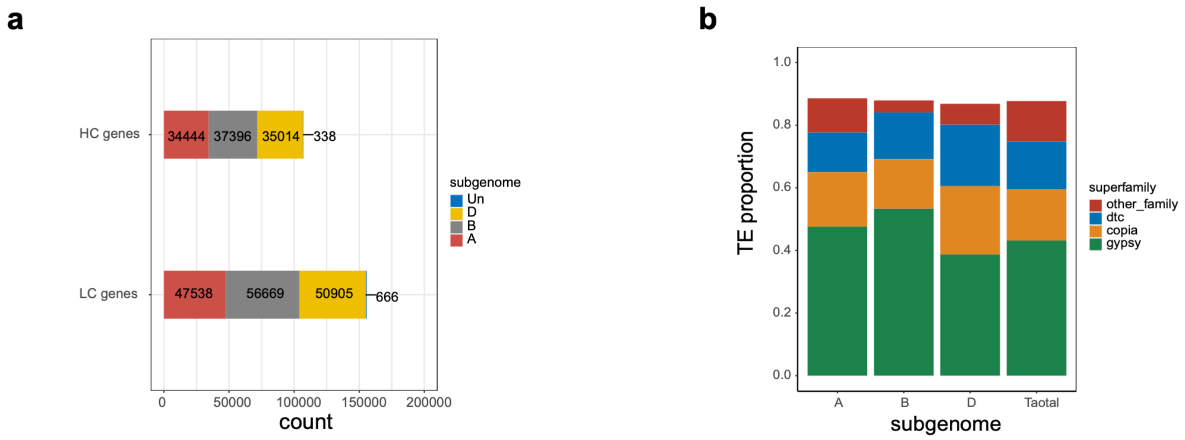


**Supplementary Fig. 7. Statistics of genes and transposable elements in the JM44 genome. a**, Selected gene prediction statistics of the JM44 genome, including number and subgenome distribution of HC and LC genes. **b**, Transposable elements statistics of the JM44 genome.

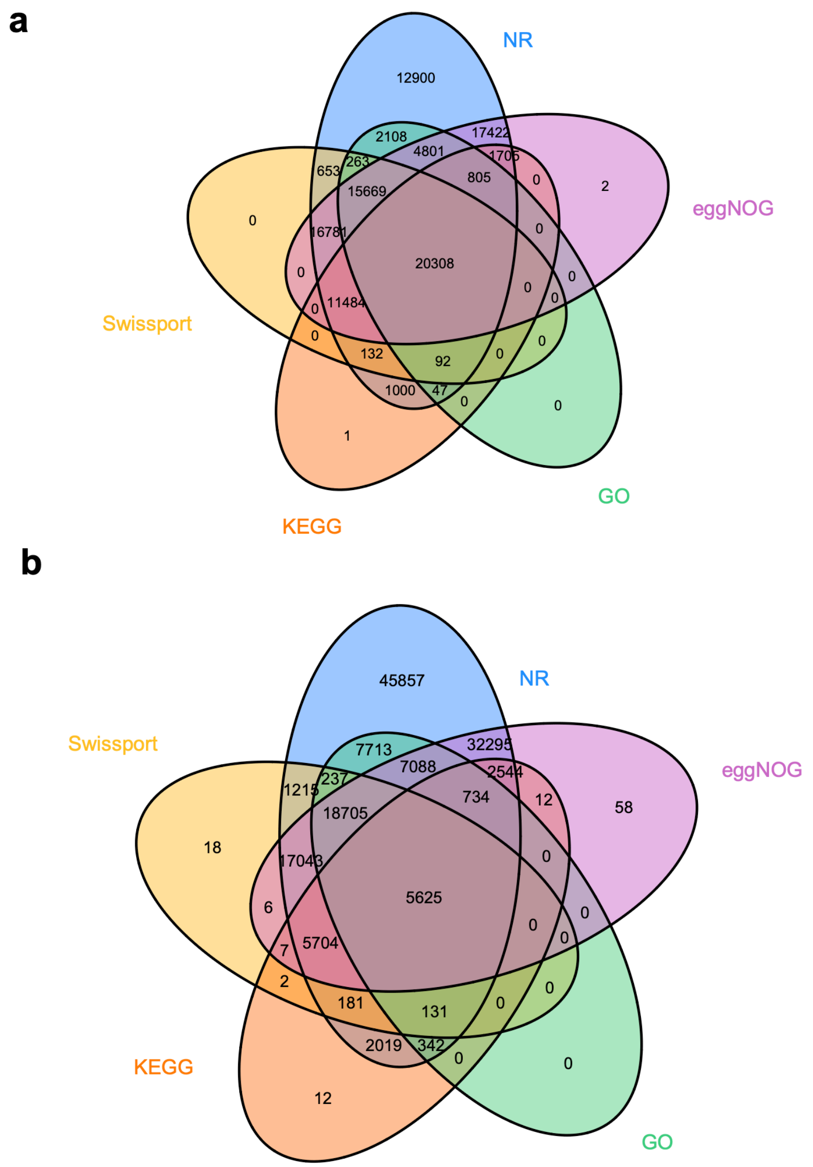


**Supplementary Fig. 8. Venn diagram of gene function annotation in different databases. a** corresponds to the HC gene set and **b** to the LC gene set.

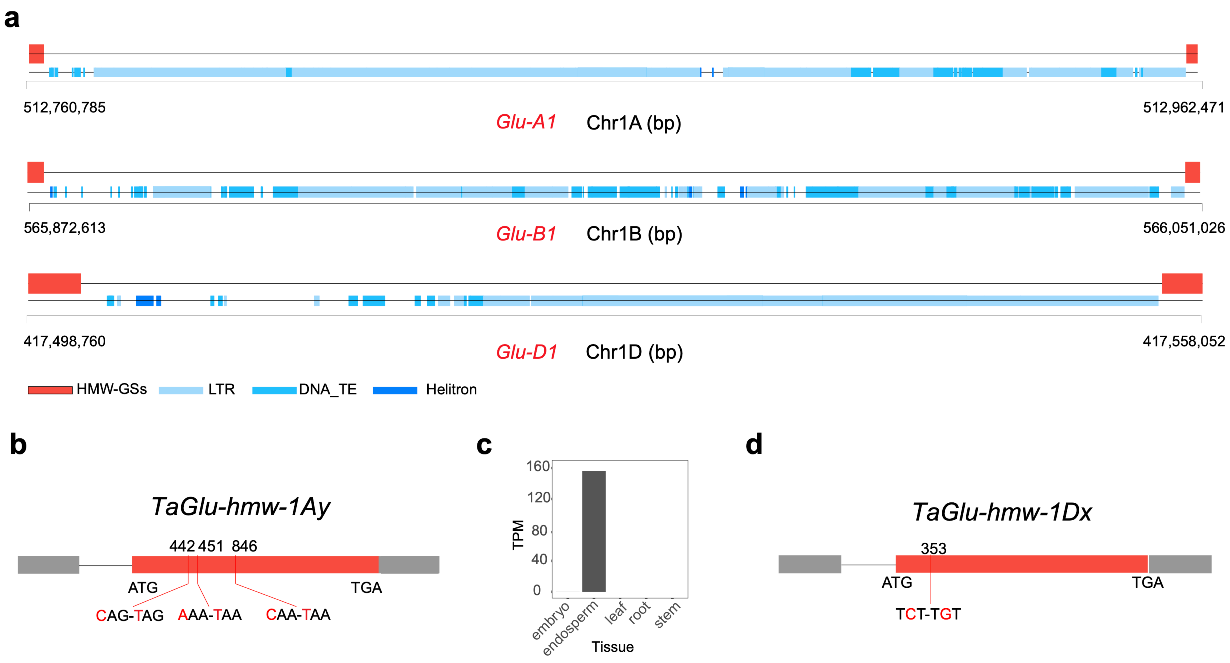


**Supplementary Fig. 9. Structural analysis of the *Glu-1* locus. a, Precise structural organization of the *Glu-1* locus. b,** Silencing-associated mutation sites within the *TaGlu-hmw-1Ay* gene. **c,** Expression levels of *TaGlu-hmw-1Ay* in root, stem, leaf, embryo, and endosperm at 15 days after pollination. **d,** Key mutation sites within the *TaGlu-hmw-1Dx* gene.

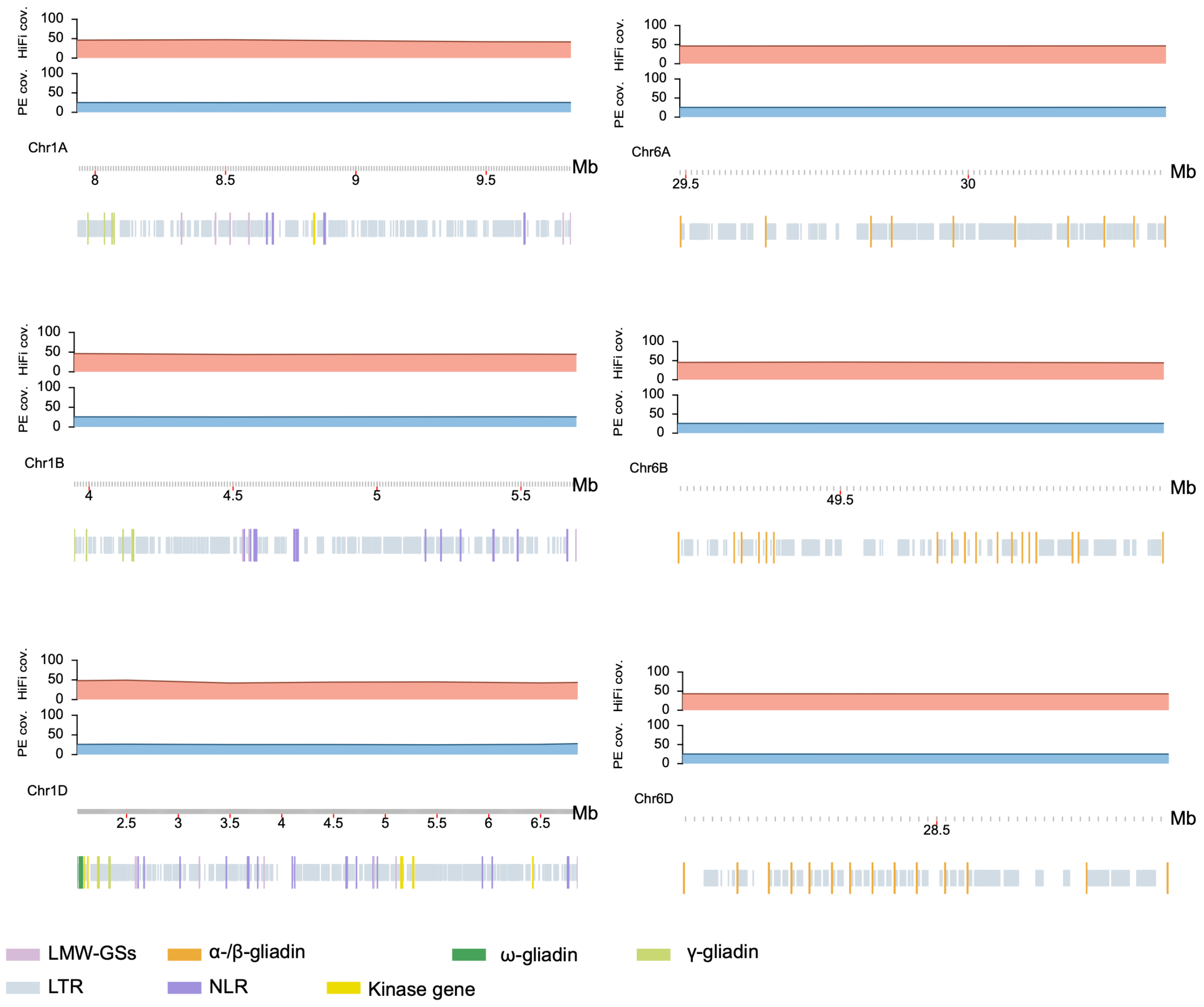


**Supplementary Fig. 10. Complete assembly of LMW-GSs and gliadin gene clusters.** The genome-wide coverage of HiFi and Illumina reads mapped are in the first and second layers from top to bottom, respectively. A uniform depth distribution indicates the complete assembly of the gluten gene loci.

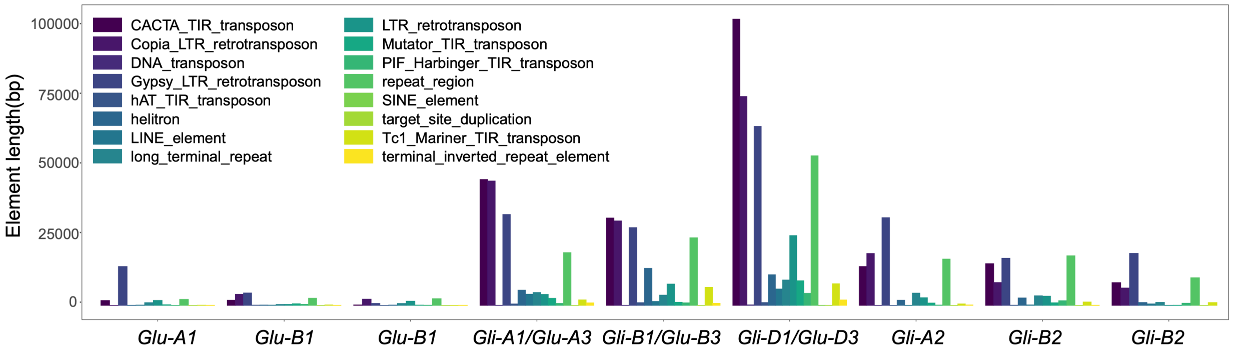


**Supplementary Fig. 11. Cumulative lengths of different types of repetitive elements within the gluten gene loci.**

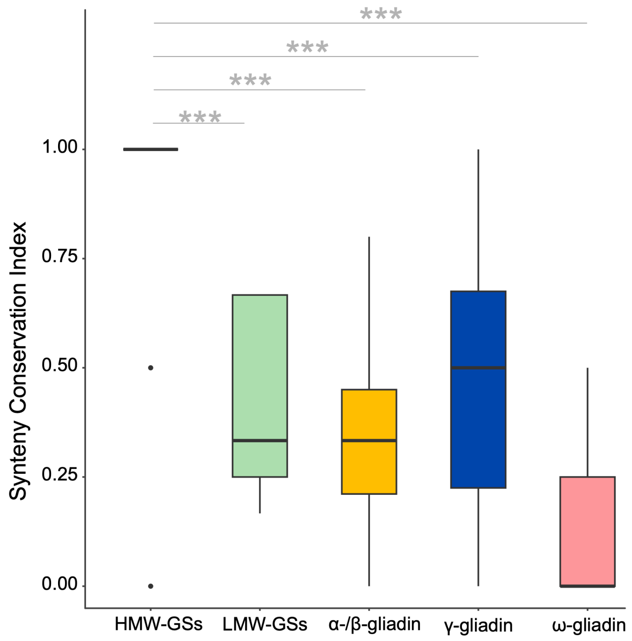


**Supplementary Fig. 12. Distribution of Synteny Conservation Index (SCI) for different types of gluten genes.** *P* values were generated using one-tailed *t*-tests.

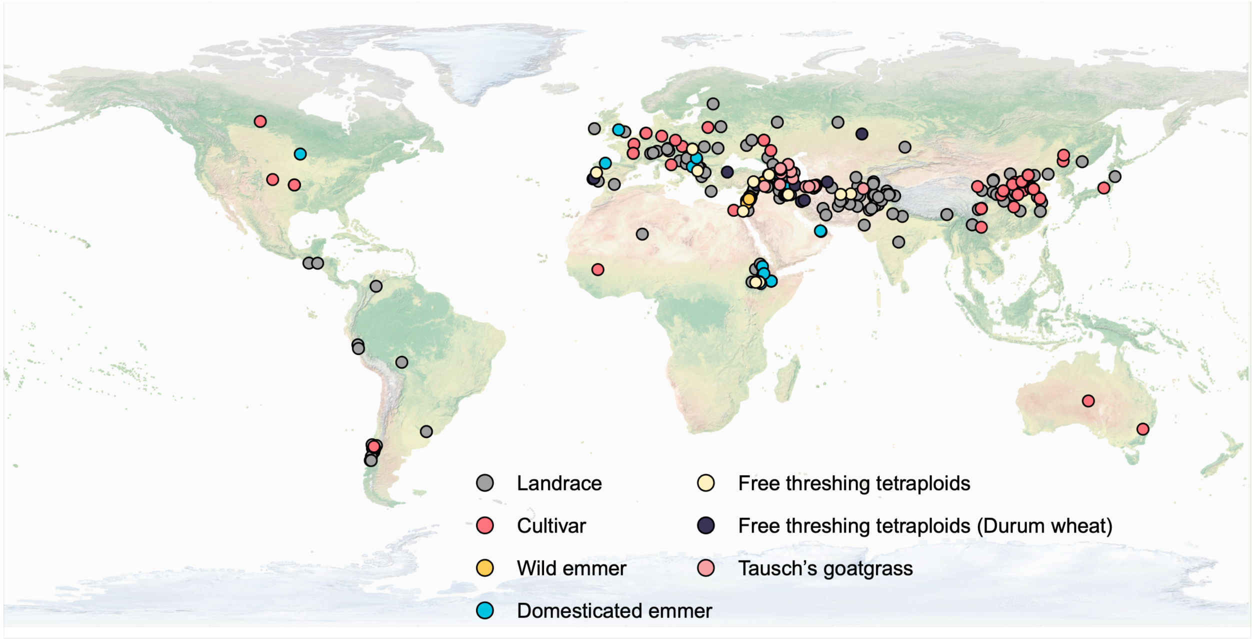


**Supplementary Fig. 13. Global sampling map of JVMap.** Different colors represent different sample categories.

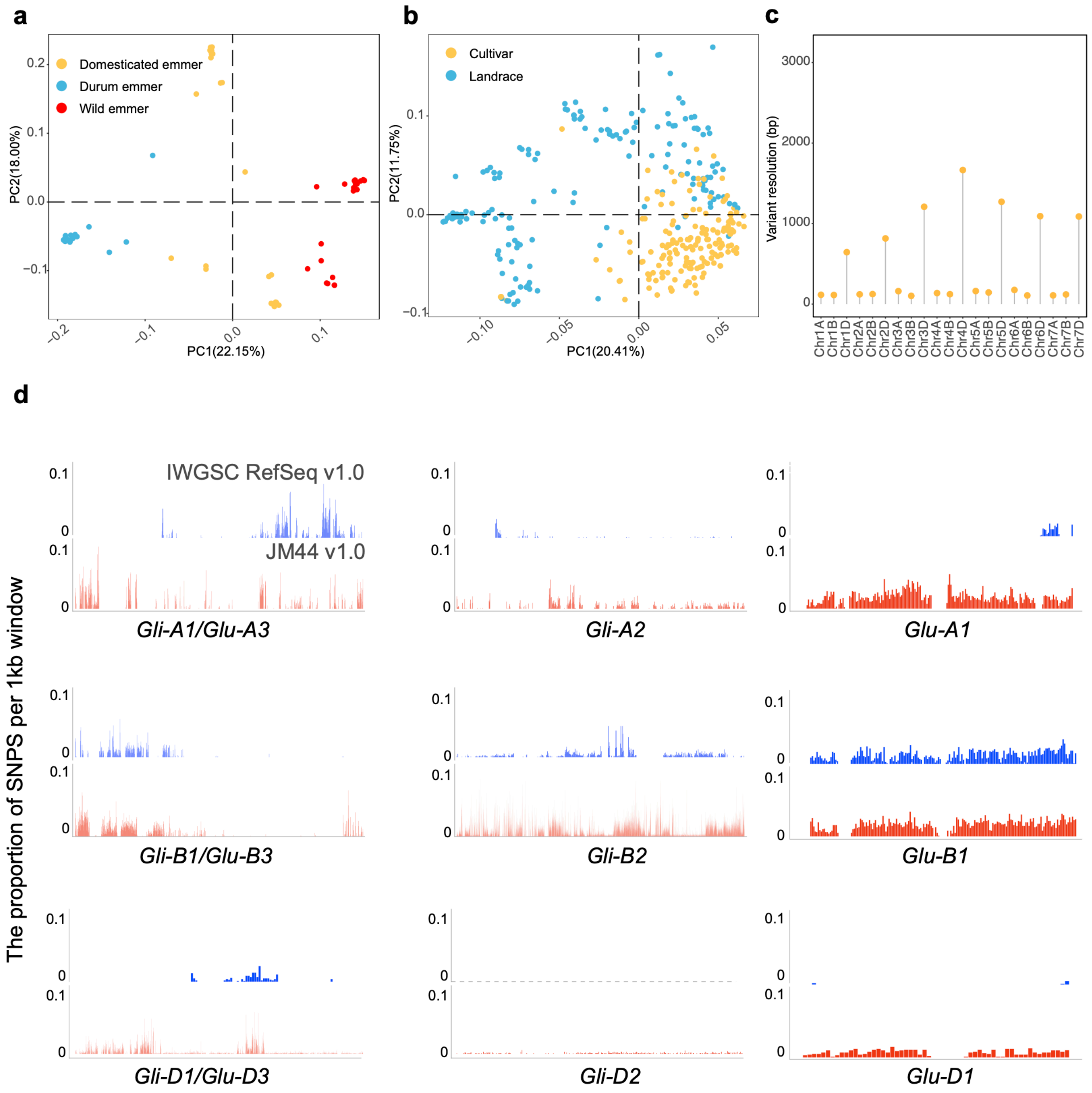


**Supplementary Fig. 14. JVMap variant overview.** **a** and **b** show principal component analysis (PCA) of tetraploid and hexaploid samples, respectively. **c** shows the variant resolution across different chromosomes. **d** shows the SNP distribution within the gluten gene loci.

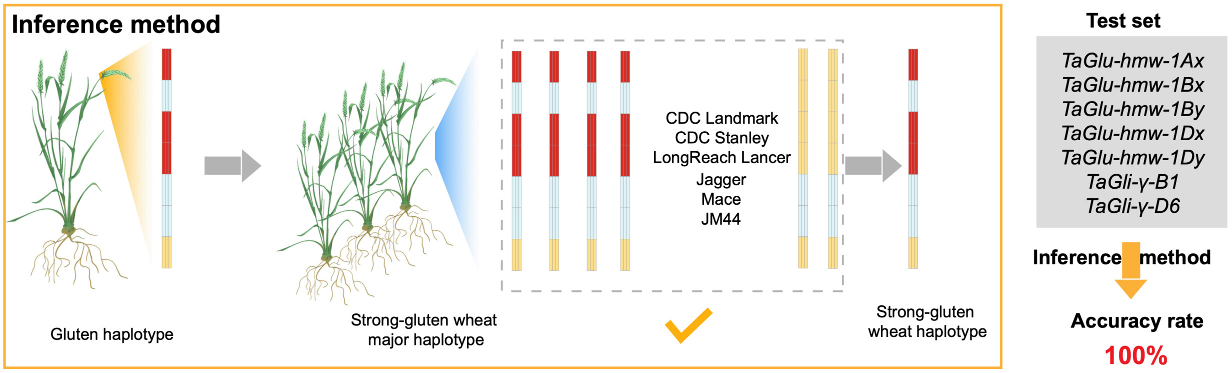


**Supplementary Fig. 15. Schematic of the inference method for strong-gluten haplotypes of gluten genes.** The genes in the test set on the right are published and confirmed to have strong-gluten haplotypes.

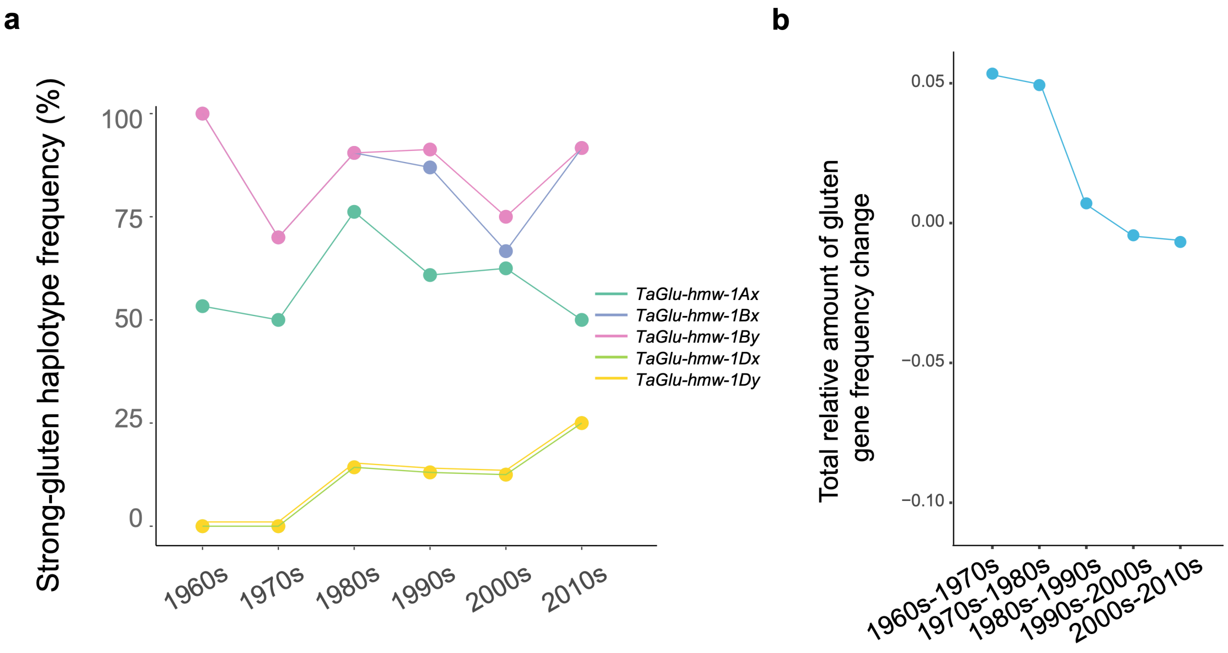


**Supplementary Fig. 16. Gluten gene variation frequencies during modern wheat breeding in China.** **a**, Changes in the frequency of strong-gluten haplotypes of HMW-GSs genes across different decades during the modern breeding in China. **b**, Relative changes in allele frequencies of all gluten genes between different decades during the modern breeding in China.

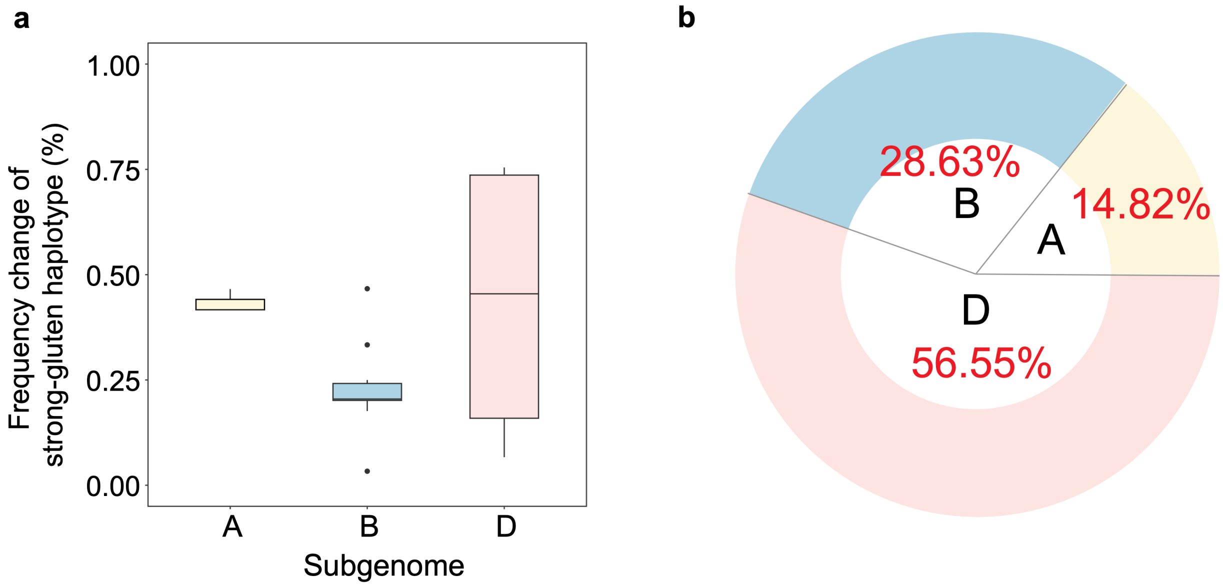


**Supplementary Fig. 17. Subgenome-level variation in strong-gluten haplotype frequencies of selected gluten genes.** **a**, Magnitude of strong-gluten haplotype frequency changes among selected gluten genes, grouped by subgenome. **b**, Relative contribution of each subgenome to the total amplitude of strong-gluten haplotype frequency changes.

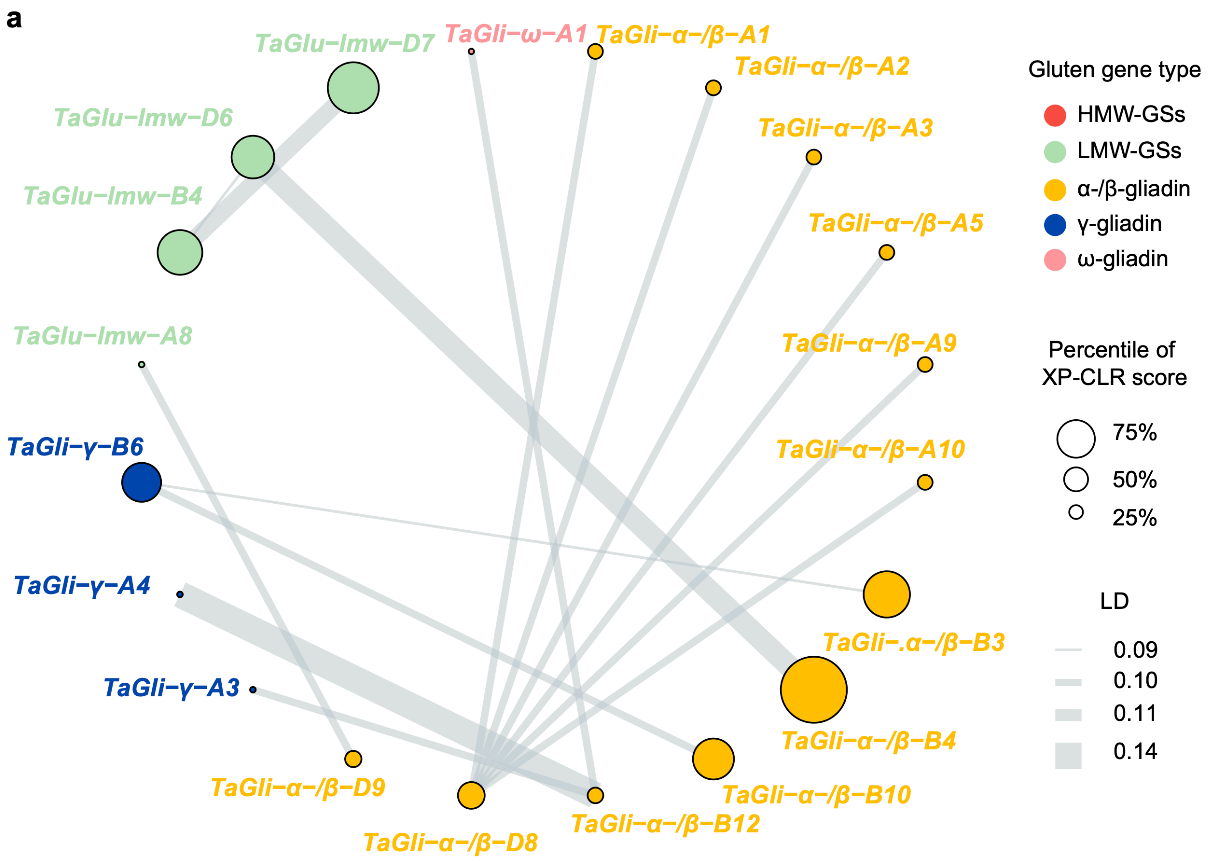


**Supplementary Fig. 18. Network of epistatic interactions among gluten genes in landraces.** Different colors represent different categories of gluten genes. The size of each gene node indicates the selection pressure experienced from landraces to Chinese modern cultivars. The thickness of each edge reflects the LD between gene pairs.

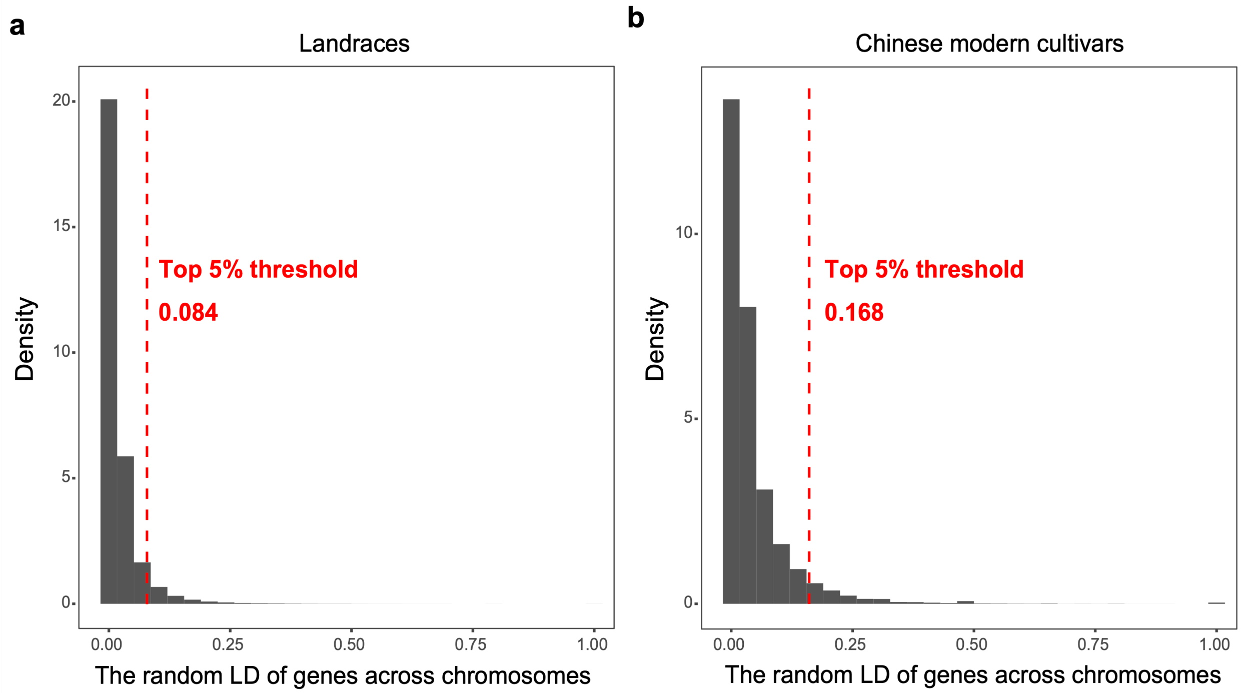


**Supplementary Fig. 19. Distribution of inter-chromosomal gene-pair LD.** **a** and **b** represent landraces and Chinese modern cultivars, respectively. The red dashed line marks the top 5% threshold at the right tail of the distribution, with the corresponding LD value indicated.

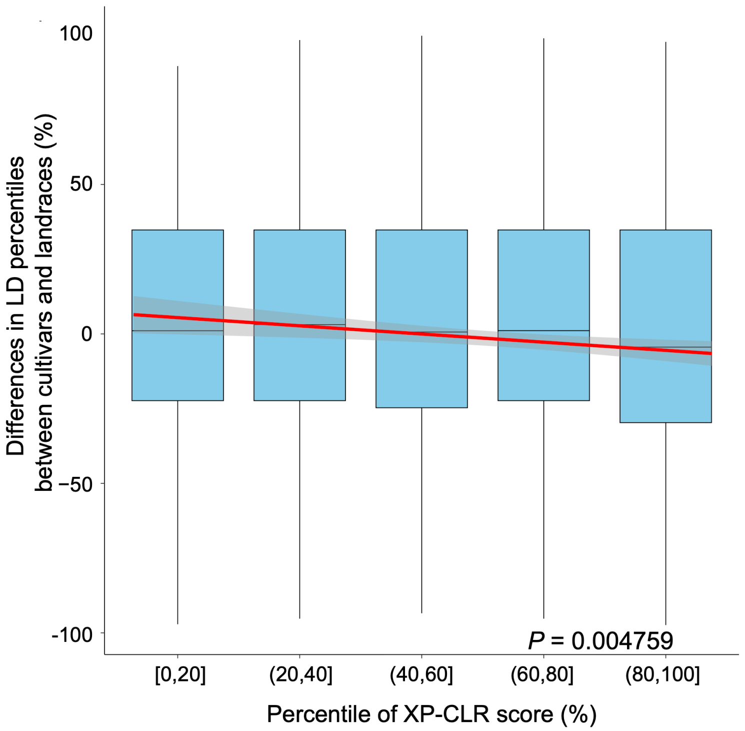


**Supplementary Fig. 20. Differences in inter-chromosomal LD percentiles of random gene loci between Chinese modern cultivars and landraces under varying selection pressures.** *P* values for fitted curves were calculated using two-tailed *t*-tests.
